# Proteomic analysis of unicellular cyanobacterium *Crocosphaera subtropica* ATCC 51142 under extended light or dark growth

**DOI:** 10.1101/2024.07.29.605499

**Authors:** Punyatoya Panda, Swagarika J. Giri, Louis Sherman, Daisuke Kihara, Uma K. Aryal

## Abstract

The daily light-dark cycle is a recurrent and predictable environmental phenomenon to which many organisms, including cyanobacteria, have evolved to adapt. Understanding how cyanobacteria alter their metabolic attributes in response to subjective light or dark growth may provide key features for developing strains with improved photosynthetic efficiency and applications in enhanced carbon sequestration and renewable energy. Here, we undertook a label-free proteomic approach to investigate the effect of extended light (LL) or extended dark (DD) conditions on the unicellular cyanobacterium *Crocosphaera subtropica* ATCC 51142. We quantified 2287 proteins, of which 603 proteins were significantly different between the two growth conditions. These proteins represent several biological processes, including photosynthetic electron transport, carbon fixation, stress responses, translation, and protein degradation. One significant observation is the regulation of over two dozen proteases, including ATP dependent Clp-proteases (endopeptidases) and metalloproteases, the majority of which were upregulated in LL compared to DD. This suggests that proteases play a crucial role in the regulation and maintenance of photosynthesis, especially the PSI and PSII components. The higher protease activity in LL indicates a need for more frequent degradation and repair of certain photosynthetic components, highlighting the dynamic nature of protein turnover and quality control mechanisms in response to prolonged light exposure. The results enhance our understanding of how *Crocosphaera subtropica* ATCC51142 adjusts its molecular machinery in response to extended light or dark growth conditions.

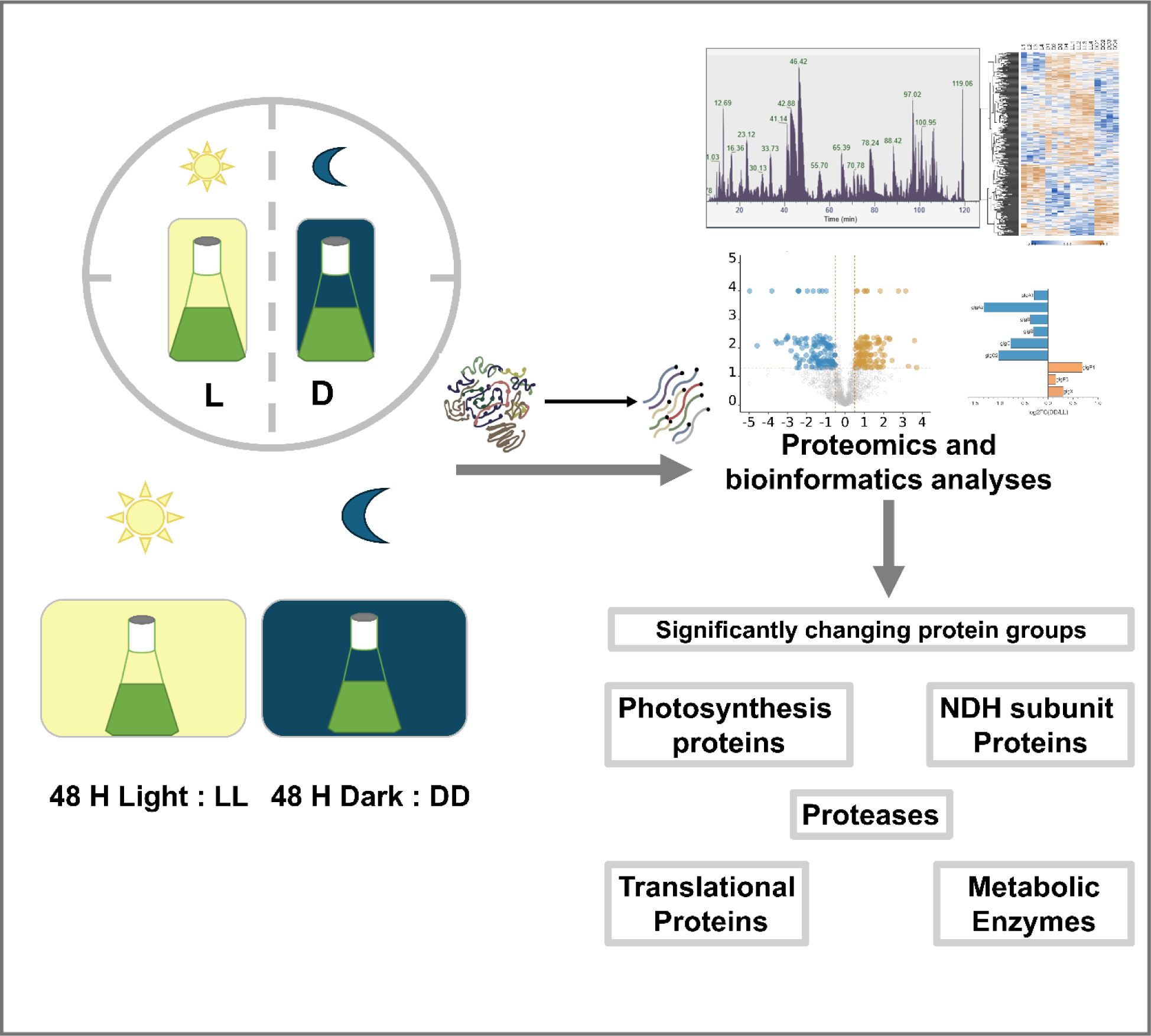

## Introduction

Cyanobacteria play an important role in harvesting solar energy and in the evolution of atmospheric oxygen. Only a few cyanobacteria including *Crocosphaera subtropica* ATCC 51142 (formerly known as *Cyanothece sp*. 51142), are capable of biological N_2_-fixation (BNF) within a single cell [1, 2]. *Crocosphaera* 51142 produces oxygen and stores photosynthetically fixed carbon in the form of glycogen granules during the day, and subsequently metabolizes stored carbon at night to produce excess energy and create an O_2_-limited intracellular environment [3, 4]. Respiratory electron transport scavenges oxygen to establish anaerobic intracellular conditions necessary for N_2_-fixation because BNF is carried out by the nitrogenase enzyme complex which is sensitive to oxygen [3, 5]. Transcriptomics [6] and proteomics [1, 2, 7-9] analyses have provided unprecedented information to understand cyanobacterial physiology and metabolism and how cells use light-dark cycles to permit the functions of two antagonistic metabolic processes.

*Crocosphaera* 51142 is a relatively well-studied model for fundamental processes of photosynthesis, respiration, CO_2_- and N_2_-fixation [10]. Cyanobacteria are also the only known prokaryotes that have evolved circadian rhythms [11]. The anticipation of impending environmental changes such as diurnal light-dark rhythms are well documented at both the transcriptional [1] and protein levels [7, 8, 12, 13]. Under normal diurnal growth cycles, their metabolic machinery is in constant dynamic change throughout the course of a day, cycling between O_2_ production in photosynthesis during the day and O_2_ scavenging during respiration and N_2_-fixation at night, with peaks every 24 hours. The accumulation and degradation of glycogen and cyanophycin (nitrogen-rich, polypeptide-like storage material) granules occur concomitantly with photosynthesis and N_2_-fixation [4, 14]. However, how their metabolic machinery responds and adjusts to subjective growth conditions such as extended light (LL) or extended dark (DD) periods compared to normal periods is not well understood.

Previous proteomic analyses of this species have primarily been carried out under predictable alternating light-dark diurnal cycles. [6, 7, 13]. However, *Crocosphaera* 51142 is also known for its circadian nature and unique metabolic capabilities such as high rates of nitrogenase-mediated H_2_ production under aerobic conditions [15-17]. However, how it responds to unpredictable environmental conditions such as extended periods of light or darkness is unknown. By subjecting cultures to 48-hour periods of continuous light (LL) or continuous darkness (DD) after several days of normal 12-hour light (L) and 12-hour dark (D) growth, we sought to investigate how this strain adapts to such conditions at the proteomic level. The observed changes in the abundances of various proteins involved in cellular processes such as metabolism, energy production, protein degradation, and protein translation reflect the organism’s adaptive mechanisms that contribute to the ecological success of the species. Understanding these adaptive responses can provide valuable insights into how organisms like *Crocosphaera* 51142 thrive in diverse environmental conditions with implication in biotechnology.

## Materials and Methods

### Culture growth Conditions

*Crocosphaera sp. subtropica* 51142 stock cultures were maintained in ASP2 medium with 17.6 mM NaNO_3_ and 50 µmol photons m^-2^ s^-1^ of continuous light. One-tenth (10ml) of the *Crocosphaera* 51142 stock was first inoculated to each 250-ml flask containing 100 mL ASP2 medium with 17.6mM NaNO_3_ and allowed to grow for 7 days on a shaker at 125 rpm, 30°C, and 50 µmol photons m^-2^ s^-1^ of continuous light. After seven days, *Crocosphaera* 51142 cultures were transitioned to 12 h-light/12 h-dark (50 µmol photons m^-2^ s^-1^) growth condition and grown for additional seven days to mid log phase. Then, cells were harvested at 4 h into the light period (L) or 4 h into the dark period (D). The remaining cultures were then grown for an additional 48 hours of either continuous light (LL) or continuous dark (DD) before harvesting.

Cells were collected by centrifugation at 3,220*g* in 15ml tubes (Corning). Cell pellets were gently washed with 1ml of 50mM HEPES-KOH buffer (pH 7.5) to remove residual growth medium, centrifuged again at 10,000*g* at 4◦C for 15 min to obtain the pellet. The pellets were resuspended in 200 μl of HEPES/KOH (pH 7.5) buffer, supplemented with 1 mM phenylmethylsulfonylfluoride (PMSF) protease inhibitor and homogenized in Precellys VK 0.5 tubes (Bertin Corp., Rockville, MD, USA), 3× at 6,000 rpm for 3 x 20 secs in each cycle followed by probe sonication. Protein concentration was determined using bicinchoninic acid (BCA) assay (Pierce Chemical Co., Rockford, IL, USA), using Bovine serum albumin as a standard and volumes corresponding to 100 μg of total protein were ultracentrifuged at 150,000 × g for 20 min to divide proteins into soluble and insoluble fractions. The soluble fractions were acetone precipitated with four volumes of ice-cold acetone and incubated overnight at -20°C, while the insoluble pellets were further resuspended with 200μl of HEPES/KOH (pH 7.5), bath sonicated for 5 mins and then precipitated with four volumes of cold (-20°C) acetone. The soluble and insoluble pellets were obtained by centrifuging at 17,200 *× g* for 20 min at 4°C, washed 3× with 80% cold (-20°C) acetone and processed for MS analysis.

### Sample preparation for LC-MS/MS analysis

Both soluble and insoluble pellets were resuspended with 23μl buffer (5% sodium dodecyl sulfate (SDS), 50 mM triethylammonium bicarbonate (TEAB; pH 8.5)) and were solubilized by vortexing for two hours. The samples were then reduced by adding dithiothreitol (DTT) to a final concentration of 10 mM and incubated at 55°C for 30 minutes. The samples were then cysteine alkylated with iodoacetamide (IAA) to a final concentration of 25 mM and incubating in the dark for 30 minutes. The reduced and alkylated samples were acidified with 2.5% phosphoric acid to completely denature the proteins, followed by the addition of 165 μl of binding/wash buffer (100 mM TEAB in 90% methanol) and mixed immediately. These were passed through the S-Trap micro spin columns (Protifi, USA) and centrifuged at 2,000 *x g* for 1 minute. The S-trap columns were washed three times with 150µl binding/wash buffer with centrifugation at 2,000 x g. These columns were then transferred to clean 1.7mL Eppendorf tubes and the trapped proteins were digested overnight with 1 μg of MS grade Pierce trypsin protease (ThermoFisher Scientific, USA) resuspended in 50mM TEAB buffer, in a thermomixer at 37°C. Purified peptides were eluted using 50% acetonitrile, 50 mM TEAB and 0.2% formic acid via centrifugation with each for 1min at 2000 x g and subsequently dried in a vacuum centrifuge (Vacufuge Plus, Eppendorf, Enfield, CT)) at 45°C.

### Reversed phase LC separation and tandem mass spectrometry analysis

Dried desalted peptides were resuspended in resuspension solution composed with 3% acetonitrile (ACN), 0.1% formic acid (FA) in water to a final concentration of 1μg/μl and 1μg of peptides were used for subsequent LC-MS/MS analysis. Peptides were analyzed by reverse-phase separation using the Dionex UltiMate 3000 RSLC nano system, coupled to a Q-Exactive High-Field (HF) Hybrid Quadrupole Orbitrap MS via Nanospray Flex^TM^ electrospray ionization source (Thermo Fisher Scientific) as described previously [18, 19]. Briefly, peptides were first loaded into a trap column (300 μm ID × 5 mm) packed with 5 μm 100 Å PepMap C18 medium (Thermo Fisher Scientific, Waltham, MA, USA), and then separated using a reverse phase 1.7 µm 120 Å IonOptics Aurora Ultimate C18 column (75 µm x, 25cm). The column was maintained at 40 °C, mobile phase solvent A was 0.1% FA in water, solvent B was 0.1% FA in 80% ACN. The loading buffer was 0.1% FA in 2% ACN. Peptides were loaded into the trap column for 5 min at 5 μl/min, then separated with a flow rate of 300 nl/min using a 130 min linear gradient. The concentration of mobile phase B was increased linearly to 8% in five minutes, 27% B in 80 min, and then 45% B at 100 min. After 100 min, it was subsequentially increased to 100% of B at 105 min and held constant for another 7 min before reverting to 2% of B in 112.1 min and maintained at 2% B until the end of the run. The QE-HF was operated in positive ion and standard data dependent acquisition (DDA) mode. The spray voltage was set at 2.6 kV, the capillary temperature was 320 °C and the S-lens RF was set at 50. The resolution of Orbitrap mass analyzer was set to 60,000 and 15,000 at 200 *m/z* for MS1 and MS2, respectively, with a maximum injection time of 100 ms for MS1 and 20 ms for MS2. The full scan MS1 spectra were collected in the mass range of 350–1600 *m/z* and the MS2 first fixed mass was 100 *m/z*. The automatic gain control (ACG) target was set to 3 × 10^6^ for MS1 and 1 × 10^5^ for MS2. The fragmentation of precursor ions was accomplished by higher energy C-trap collision dissociation (HCD) at a normalized collision energy setting of 27% and an isolation window of 1.2 *m/z*. The DDA settings were for top 20 MS2 with a minimum intensity threshold of 5 × 10^4^ and a minimum AGC target of 1 × 10^3^. The dynamic exclusion was set at 15 s and accepted charge states were selected from 2 to 7 with 2 as a default charge. The exclude isotope function was activated.

### Data processing and statistical analysis

LC–MS/MS data were processed with MaxQuant analysis software (Ver 2.0.3.0) [20, 21]. Raw spectra were searched against the *Crocosphaera* 51142 protein sequence database obtained from UniProt (downloaded in July 2022) containing 5403 protein sequences, for protein identification and MS1 based label-free quantitation. The minimum length of the peptides was set at 6 AA residues in the database search. The following parameters were edited for the searches: precursor mass tolerance was set at 10 ppm, MS/MS mass tolerance was set at 20 ppm, enzyme specificity of trypsin/Lys-C enzyme allowing up to 2 missed cleavages, oxidation of methionine (M) as a variable modification and carbamidomethylation of cysteine as a fixed modification. The false discovery rate (FDR) of peptide spectral match (PSM) and protein identification was set to 0.01. The unique plus razor peptides (non-redundant, non-unique peptides assigned to the protein group with most other peptides) were used for peptide quantitation. Only proteins detected with at least one unique peptide and MS/MS ≥ 2 (spectral counts) were considered valid identifications. Label-free quantitation (LFQ) intensity values were used for relative protein abundance comparisons. The MaxQuant results were analyzed using Perseus [22] for filtering and downstream statistical analysis. Either significantly up- or downregulated proteins between the two treatment groups (L vs D and LL vs DD) were determined by a two -tailed student’s *t*-test. Differentially expressed proteins were determined with *p*-value ≤ 0.05.

### Gene Ontology (GO) assignment by ensemble of three function prediction methods

We used three sequence-based protein function prediction methods, named Protein Function Prediction (PFP) [23, 24], Phylo-PFP [25], and Extended Similarity Group (ESG) [26] to assign Gene Ontology (GO) terms to protein-coding genes. The PFP algorithm uses a scoring method based on Expect (E)-values to combine GO terms associated with sequence hits identified by PSI-BLAST Additionally, based on validation results over a set of benchmark sequences, it assigns a confidence score to GO term predictions. Phylo-PFP additionally incorporates phylogenetic information in assessing the sequence distance. The ESG method performs iterative sequence database searches and annotates a query sequence with GO terms. Each annotation is given a probability based on how similar it is to other sequences in the protein similarity graph. To ensure the inclusion of meaningful GO term annotations, we focused exclusively on predictions characterized by high confidence. Specifically, we applied confidence score cutoffs of 20,000 for PFP, 0.7 for Phylo-PFP, and 0.7 for ESG. All GO terms surpassing these respective cutoff values were incorporated into our analysis. To enhance the coverage of our predictions and include all high-confident predicted GO terms, we combined results from three prediction methods by taking the union of them and presented the consolidated GO term annotations.

Each result file has information about Protein ID, GO ID, Depth (depth of the GO ID in the GO DAG), Class (GO functional category (f-molecular function, p- Biological process, c- Cellular Component), and GO Description. The Gene Ontology release 2021-11-16 was used for this analysis.

## Results

### Overview of the proteome under different growth conditions

The focus of this study was to determine changes in the *Crocosphaera* 51142 proteome during growth under extended light or dark conditions. However, for comparative purposes, we also analyzed samples grown under normal 12h L/D cycles (Figure 1). We identified 24,491 peptides (Supplementary Table S1) and 2287 proteins (Supplementary Table S2) at 1% FDR. After further filtering of proteins to retain those with LFQ values > 0 and MS/MS counts ≥ 2 in at least three of the four replicates in at least one treatment group, we retained 1640 (70%) proteins with appropriate quantitative values for statistical comparison (Supplementary Table S3). The Venn diagram (Figure 2A) shows the overlap of proteins identified in each group. 1305 proteins were identified in L/D growth and 1585 proteins were found in LL/DD. Principal Component Analysis (PCA) of relative intensities of all proteins revealed a clear separation among the four treatment groups, indicating distinct proteomic profiles associated with each experimental condition with 32.7% explained variance (Figure 2B). Furthermore, the PCA plot demonstrates that replicates within each treatment group were clustered together, indicating the biological reproducibility of the samples. The heat map (Supplementary Figure 1A) illustrates the qualitative and quantitative differences of protein candidates across all samples.

**Figure 1.**
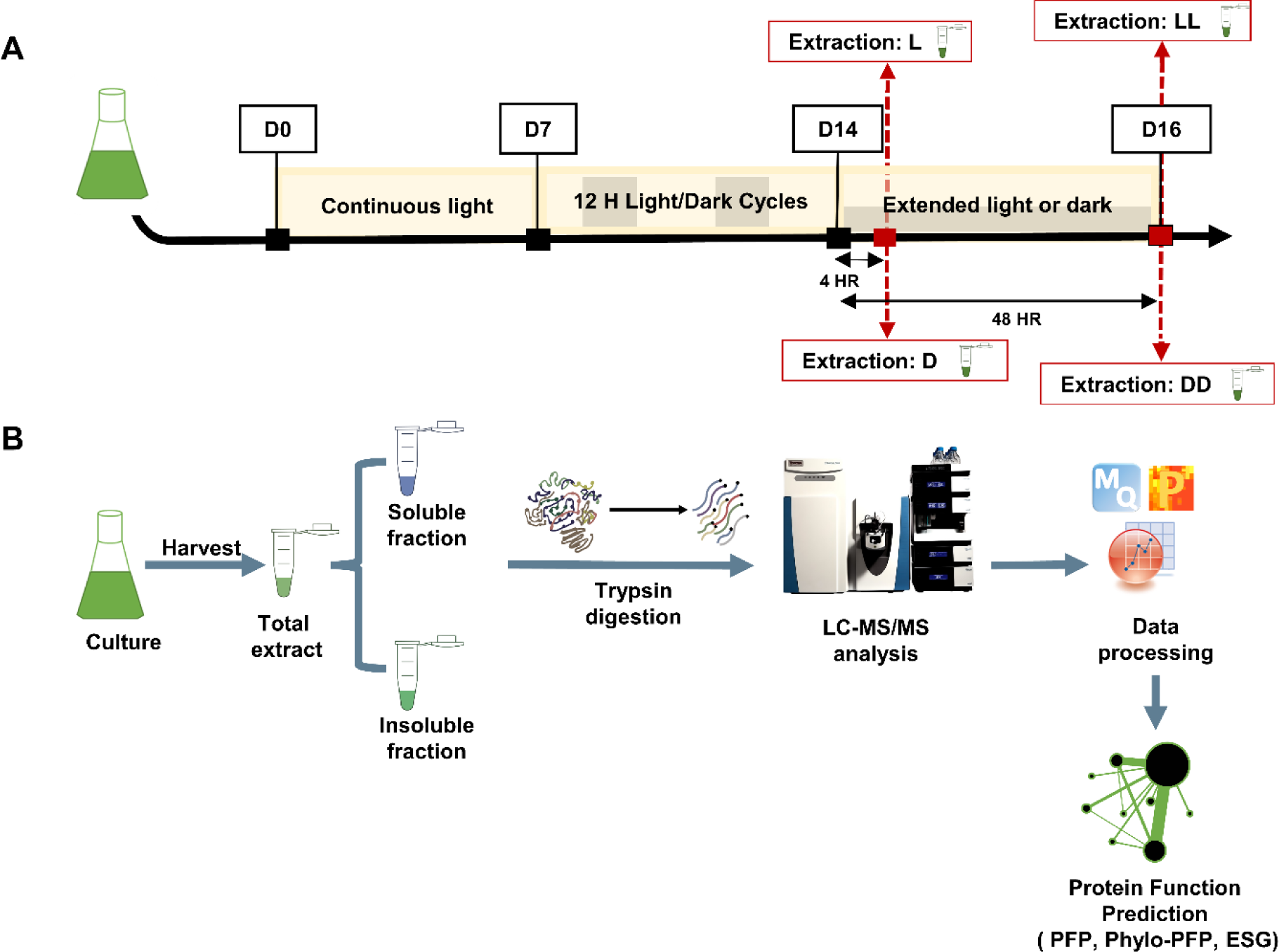
Experimental design and proteomics workflow. **(A).** *Crocosphaera subtropica* cultures were grown in ASP2 medium supplemented with NaNO_3_ at 30°C under continuous light for seven days. Cells were then transitioned to 12-hour light-dark cycles and grown for seven more days before harvesting. Fifty (50) ml of growing cell cultures were harvested at four hours into the light (L) or 4 hours into the dark (D). The remaining cultures grew for 48 more hours at either continuous light (LL) or continuous dark (DD) before harvesting. **(B).** Cell lysates from four biological replicates were divided into soluble and insoluble fractions, digested with trypsin, and analyzed by LC-MS/MS. Data processing was done using MaxQuant, followed by statistical analysis using Perseus and OriginPro. Function prediction was done using three sequence-based prediction methods, Protein Function Prediction (PFP), Phylo-PFP and Extended Similarity Group (ESG), and the results from all the three methods were combined to assign GO terms to protein- coding genes.

**Figure 2.**
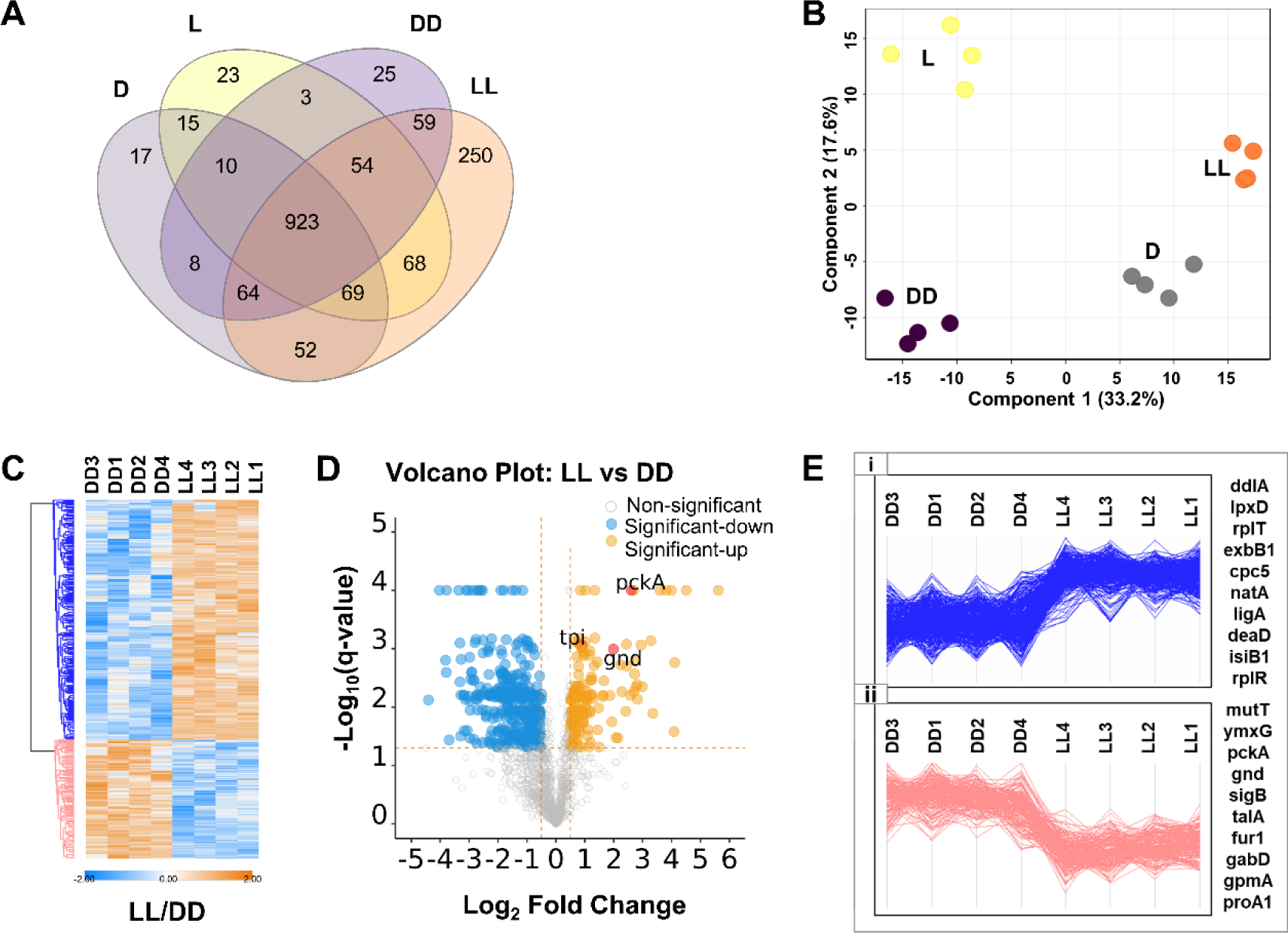
Overview of *Crocosphaera* 51142 proteomes during growth under light-dark diurnal cycles and extended light (LL) or extended dark (DD). **(A).** Venn Diagram showing the number of proteins overlapping among different growth conditions. **(B).** Principal Component Analysis of all the proteins identified across light (L), dark (D), extended light (LL) and extended dark (DD) samples. **(C).** Heatmaps of 603 significantly changing proteins between extended light and extended dark conditions. **(D).** Volcano plot of all quantified proteins under extended light and extended dark conditions. The horizontal line represents the *log_2_* (*fold change)* cutoff*, and the* vertical line represents the *-log_10_ (q-value)* cutoff. (E). Cluster profiles of significant proteins. The top 10 proteins with the highest and lowest fold change in LL and DD conditions are indicated on the right side of figure E.

We performed a student’s *t*-test to compare proteins that were differentially abundant between treatment conditions and set the *q*-value at < 0.05 [18, 19, 27, 28], as the minimum cutoff to consider proteins as “significantly regulated”. Using this cut-off, we identified 315 proteins significant between L/D (Supplementary Table S4) and 603 significant between LL/DD (Supplementary Table S5). A heatmap representation of all the significant proteins demonstrates that the distribution of upregulated and downregulated proteins was symmetrical in L/D samples (Supplementary Figure 1B). However, there were more proteins that were upregulated in LL than in DD (Figure 2C). The volcano plot (Figure 2D, Supplementary Figure 1C) shows distribution of up- and down-regulated protein in LL compared to DD. The distribution of abundance patterns of significant proteins between LL and DD are displayed in Figure 2E and Supplementary Figure 1D. The gene symbols on the right of Figure 2E show the top 10 most up- and down-regulated proteins in LL compared to DD. Adenine DNA glycosylase (MutT), processing protease (YmxG), phosphoenolpyruvate carboxykinase (ATP) (Pcka), 6-phosphogluconate dehydrogenase (GnD), transaldolase (TalA) were among the highly upregulated proteins during extended dark. Proteins such as glycogen synthase (GlgA1), ferredoxin-dependent glutamate synthase (GlsF), allophycocyanin-B(ApcD) were the highly upregulated proteins in the extended light.

We then mapped all significant proteins to their biological functions based on UniProt and Gene Ontology (GO) terms using three different function prediction methods and the top 10 representative biological processes are shown in Figure 3. Thylakoid membrane, chloroplast and photosynthetic membranes, and ribosome were among the most represented GO-terms among upregulated proteins (Figure 3A) and structural development, cell wall macromolecule biosynthetic, and glycan biosynthetic processes were among the most represented GO-terms among the downregulated proteins between L/D (Figure 3B). Similarly, between LL and DD, NADP binding, arylsulfatase activity, vitamin binding, and nucleoside phosphate binding were the most represented GO-terms among the upregulated proteins (Figure 3C). These results suggest a potential adaptation of cells to the continuous light environment, possibly involving increased demand for NADP, an important cofactor involved in various metabolic reactions, including photosynthesis as well as increased demand for certain vitamins, sulfur, and nucleoside phosphate for energy metabolism. In contrast, glycosaminoglycan biosynthetic process, cell morphogenesis, regulation of developmental process, and cellular component organization were the most represented GO-terms among the downregulated proteins (Figure 3D), suggesting that continuous conditions may have a suppressive effect on developmental pathways, disruption of membrane organization, cell share, and structure.

**Figure 3.**
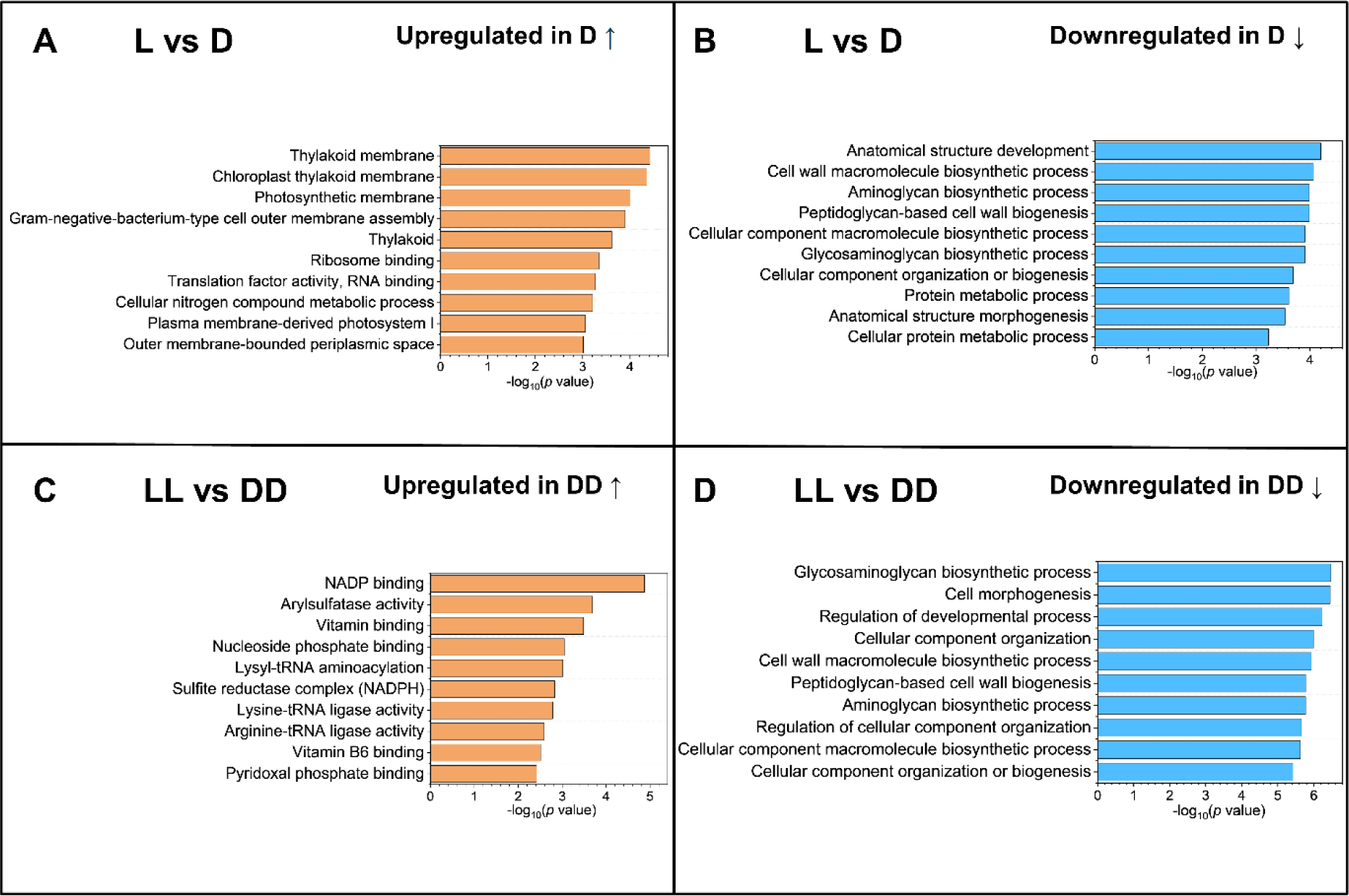
Top 10 enriched Gene Ontology (GO) terms. **(A)** Bar plots showing representative GO terms of proteins upregulated in the dark compared to the light cycle. **(B).** Bar plots showing representative GO terms of proteins downregulated in the dark compared to the light. **(C).** Bar plots showing representative GO terms of proteins upregulated in the extended dark compared to the extended light growth. **(D).** Bar plots show representative GO terms of proteins downregulated in the extended dark compared to the extended light. The –log_10_(P-values) are indicated in the X-axis.

### PSI and PSII proteins respond to extended light and dark growth

The responses of PSI and PSII proteins to light-dark conditions, especially in LL and DD, are fundamental to the understanding of how light availability influences the abundance and activity of key photosynthetic proteins. PSI and PSII proteins showed mixed responses to both L/D and LL/DD growth (Supplementary Figure 2A and 2B). Interestingly, more PSI and PSII proteins were upregulated in the dark than in the light under L/D growth condition; however, under LL and DD, more proteins were upregulated in the LL than in the DD (Table 1). The changes, however, were subtle. The upregulation of the ferredoxin-NADPH oxidoreductase (PetH, cce_2966), as well as the extrinsic cytochrome c-550 protein PsbV (cce_2955) involved in stabilizing the manganese cluster of the oxygen evolving complex [29] in the light period (Table 1) corroborate previous results [13]. The PSII subunits PsbH (cce_0860) and both the isoforms of Psb28 (cce_1599, cce_2792) were previously shown to be more abundant in the dark [13], corroborating the current results (Supplementary Figure 2A).

**Table 1.**
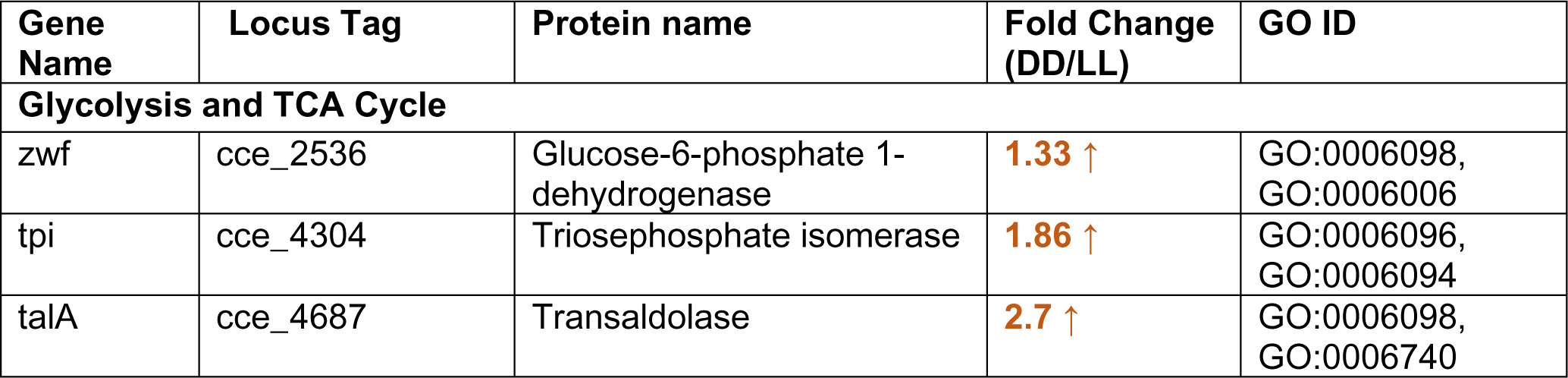
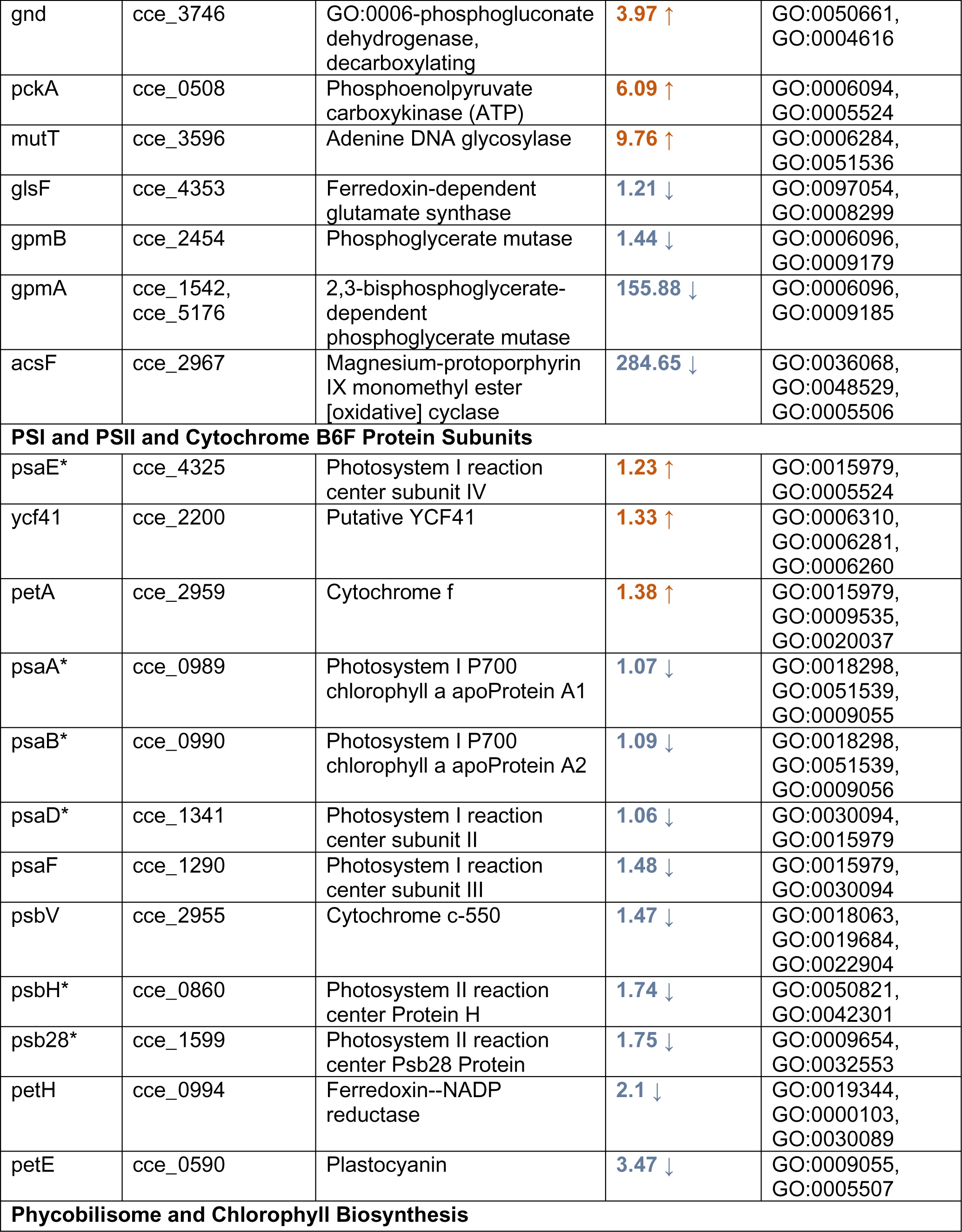
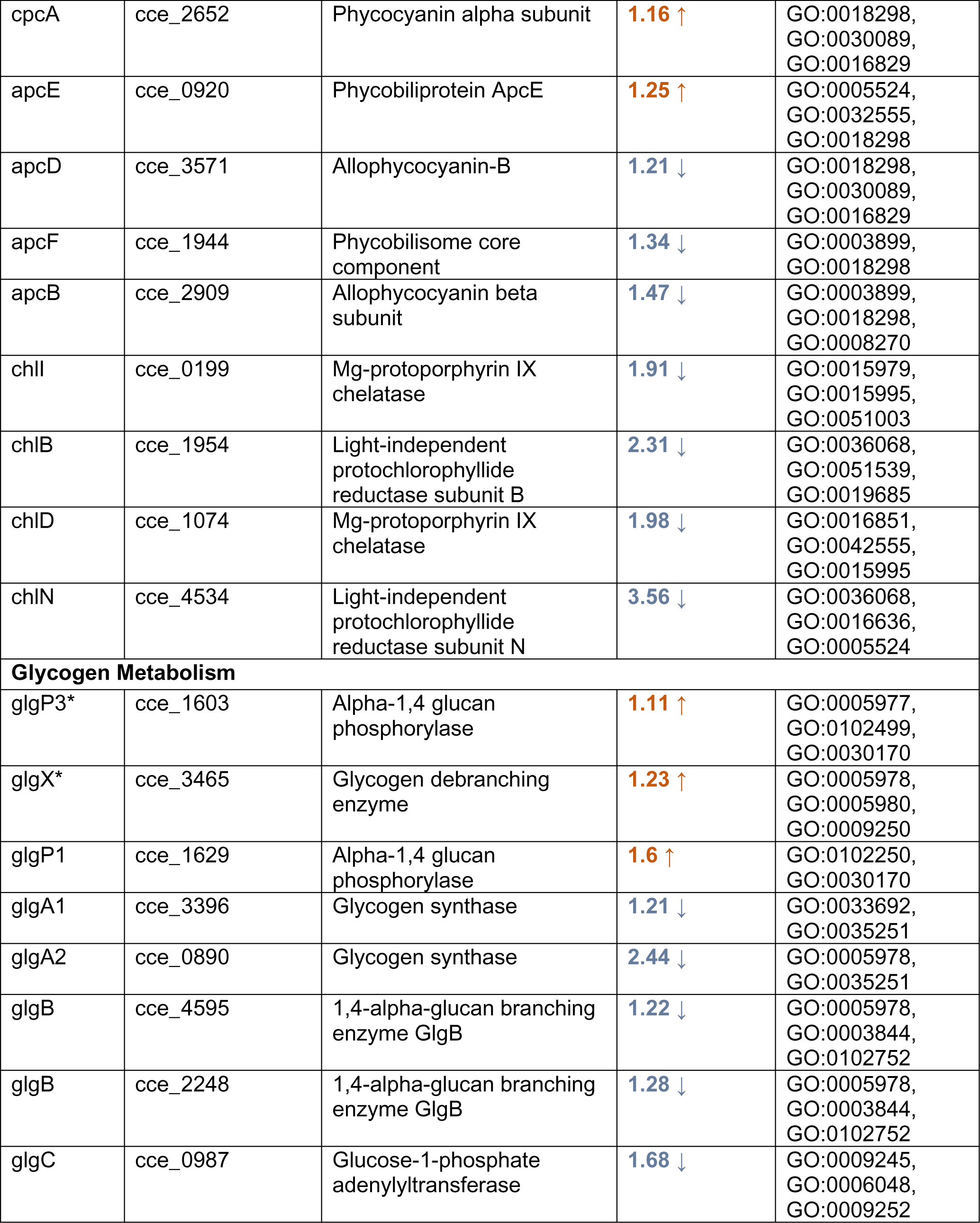
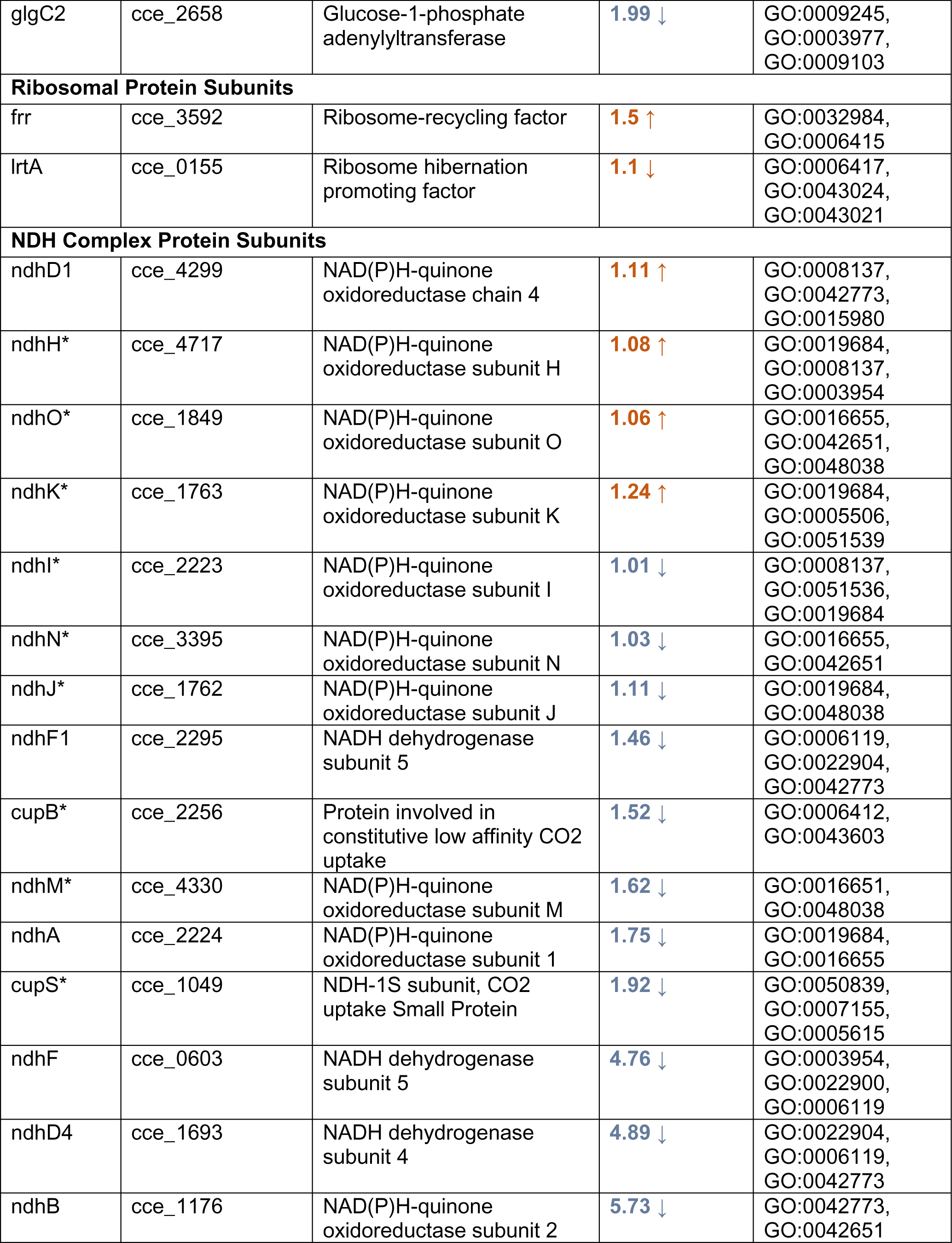
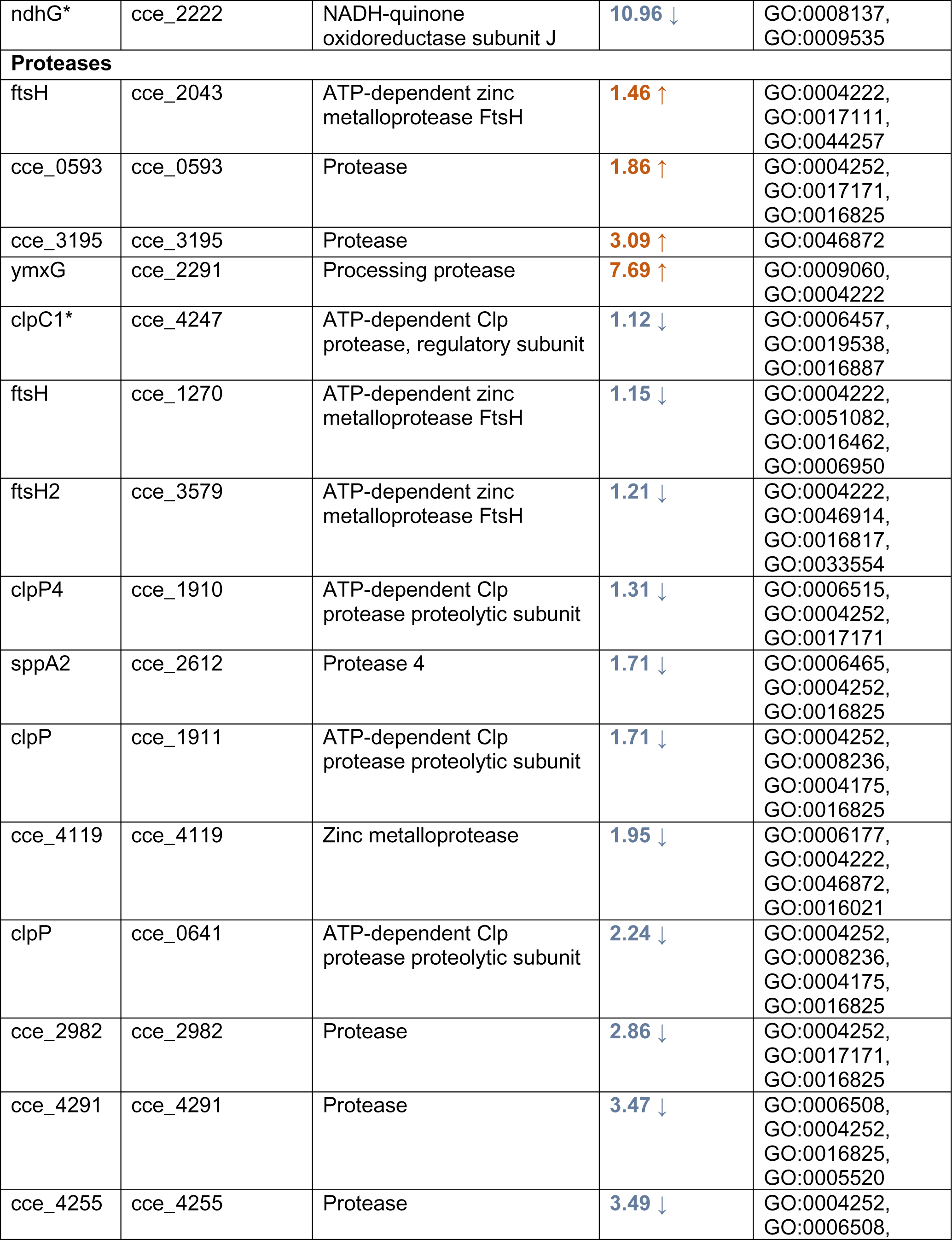
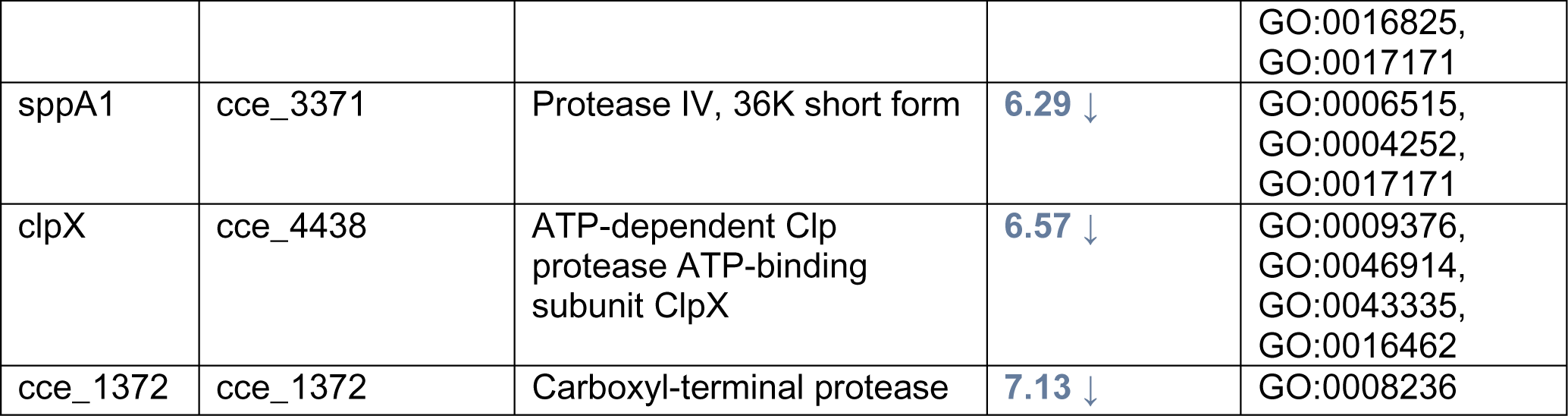
List of differentially abundant proteins identified between extended light (LL) or extended dark (DD) growth conditions along with their fold changes and associated GO terms. The numbers in orange indicate upregulation and in blue indicate downregulation in DD compared to LL (DD/LL). Proteins marked with an asterisk (*) were not significantly changing between LL or DD.

There were differences in the responses of phycobilisome (PBS) and chlorophyll biosynthesis proteins to L/D and LL/DD (Table 1). Particularly, PetE, PetH, ChlB, ChlD, ChlI, and ChlN were all more abundant in LL than in DD whereas PetA and Ycf41 were more abundant in DD than in LL (Table 1). The ApcB, ApcD, and ApcF were upregulated in the LL, but ApcE was upregulated in the DD (Table 1). Higher abundances of several photosynthetic proteins during the dark period under L/D growth, despite lower photosynthetic activity, is interesting and may indicate additional factors at play [30] which need further investigation. However, most of the PSI and PSII proteins showed subtle changes which aligns with the understanding that intrinsic membrane proteins often display more stable expression patterns compared to soluble proteins in response to environmental conditions. It may be cost effective for cells to modulate a few soluble proteins than many expensive integral membrane proteins. PSI and PSII proteins are integral components of the photosynthetic membranes, and the lipid rich membrane environment in which these proteins reside, may provide stable scaffold for their relatively stable expression, to maintain photosynthetic efficiency even under fluctuating growth conditions. The complex expression patterns of these photosynthesis proteins may also indicate their regulation by transcriptional, post-transcriptional, and post-translational mechanisms to control and modify the photosynthetic capacity and photosynthetic electron transport [13, 30, 31].

### Proteins associated with glycogen and central carbon metabolism are responsive to light- dark growth

Several rate limiting enzymes such as glucose-6-phosphate dehydrogenase (Zwf, cce_2536), transaldolase (TalA, cce_4687) and 6-phosphogluconate dehydrogenase (Gnd, cce_3746), which are involved in the OPP, had increased levels in both the dark and extended dark, suggesting increased utilization of the OPP pathway for glucose catabolism in the dark [32] to produce NADPH, an essential catabolic route for respiration. Gnd was overexpressed during the early dark periods and is primarily involved in fermentation [33, 34]. However, there are no previous reports of Gnd enzyme activity during the extended dark. This enzyme was ∼2-fold more abundant in DD than in LL, but it was only ∼1.3-fold higher in D than in L under L/D growth (Figure 2C). The higher abundance of Gnd in the DD than in the D may suggest an increased fermentation in DD to provide additional ATP to meet the energy demand of the cell.

The glycolytic enzyme, triosephosphate isomerase (Tpi, cce_4304) was also upregulated in the DD compared to LL with >2-fold increase (Table 1). Other glycolytic enzymes such as 2,3- bisphosphoglycerate-dependent phosphoglycerate mutase (GpmA, cce_1542) was also higher in DD than in LL with 2-fold increase, whereas the GpmB (cce_2454), was downregulated (Table 1). The phosphoenolpyruvate carboxykinase (PckA, cce_0508), an enzyme involved in the gluconeogenic conversion of oxaloacetate to phosphoenolpyruvate is a rare enzyme in cyanobacteria and is found only in a few species including some strains of *Crocosphaera* [2, 15]. The PckA was 6-fold higher in DD than in LL and is an important enzyme for the first step of gluconeogenesis (Figure 2D, Table 1). This enzyme would appear to be important for overall metabolism during extended darkness or other starvation conditions.

Enzymes involved in the synthesis and degradation of glycogen exhibited different responses to light-dark conditions. Glycogen synthase, GlgA1 (cce_3396) and GlgA2 (cce_0890), glycogen branching enzyme, GlgB (cce_4595), ADP-glucose pyrophosphorylase, GlgC (cce_0987) and GlgC2 (cce_2658), were all downregulated whereas glycogen phosphorylase, GlgP1 (cce_1629) was upregulated in the DD (Table 1). GlgP3 (cce_1603), another glycogen phosphorylase and glycogen debranching enzyme, GlgX (cce_3465) were also up in the DD but were not statistically significant and may suggest different forms of debranching patterns of polysaccharides exist in *Crocosphaera* 51142 to render energy for catabolic processes [35]. In *Synechocystis* sp. PCC 6803, *glgX* mutant was enriched with very short polysaccharide chains, but when mutant was transformed with *glgX* gene, the short chains decreased compared to mutant strain [36]. Interestingly, *glgX* deficient *Crocosphaera* 51142 strain maintains a greater glycogen pool and enhances dinitrogen fixation [35], thus directly linking GlgX activity with dinitrogen fixation.

### Proteases responsive to extended light- dark periods

Proteases are important in maintaining protein homeostasis, protein quality control, and recycling of damaged or misfolding proteins [37]. Unfortunately, studies focusing on proteases in cyanobacteria are limited compared to research on other aspects of cyanobacterial biology such as photosynthesis, nitrogen fixation, and ecological adaptation. Thus, the role of proteases in *Crocosphaera* 51142 and other unicellular diazotrophic cyanobacteria is an emerging area of research. We identified 29 differentially regulated proteases between L/D diurnal and extended light or extended dark, including ATP-dependent proteases (13), zinc metalloproteases (3), carboxy-terminal proteases (2), protease IV (SppA1 and SppA2), and carboxy-terminal processing CAAX protease (1). WE also identified 10 uncharacterized proteases (Figure 4, Table 1) of which seven were significantly changing between LL and DD. These observations highlight the complexity or proteolytic regulation in this organism. This dataset may represent the largest number of differentially regulated proteases highlighted in cyanobacterial research.

**Figure 4.**
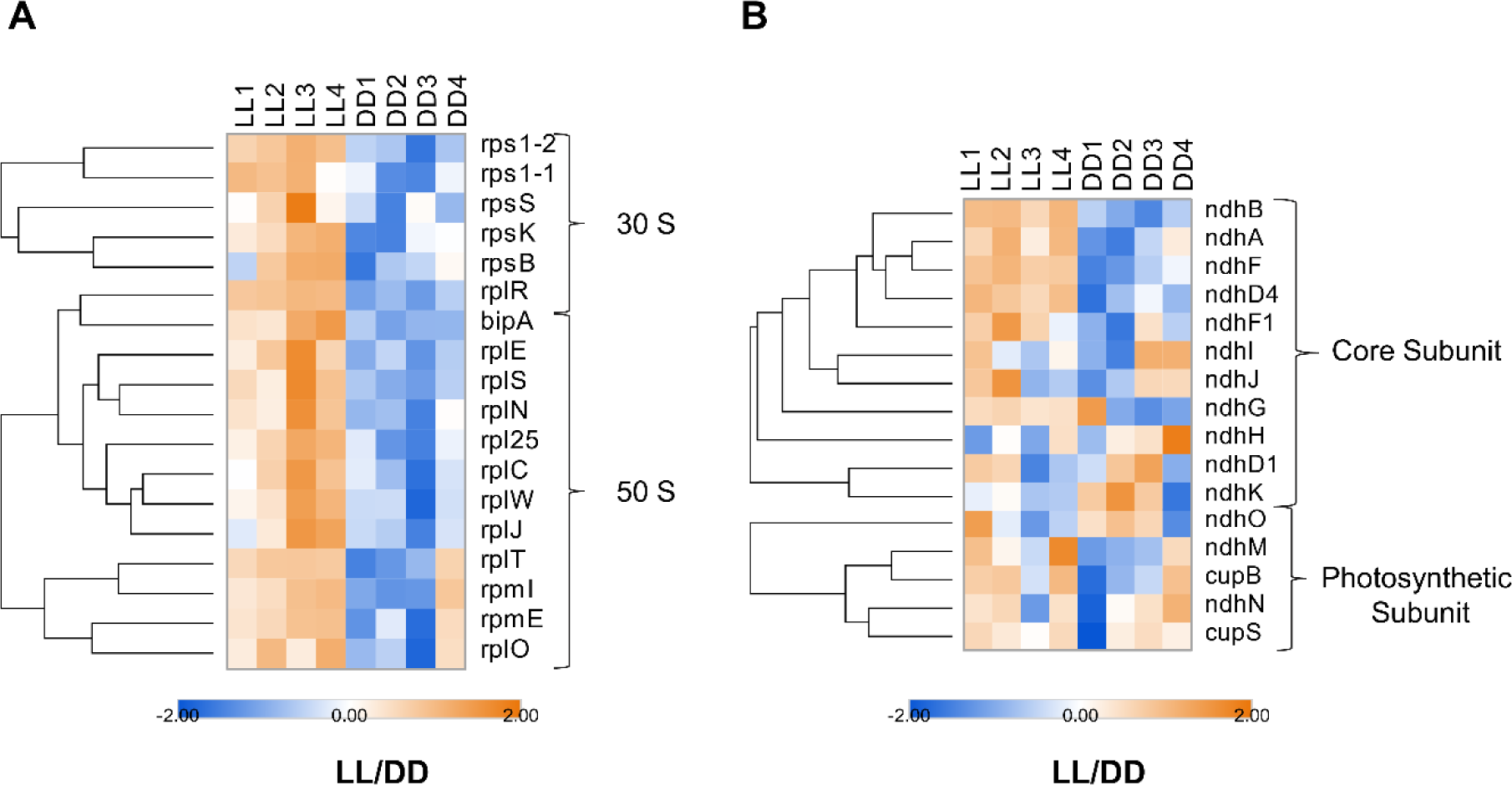
Response of different classes of proteases to different light conditions. Heatmaps depicting the z-scored log_2_(LFQ) values of differentially regulated proteases between light-dark **(A)** and extended light and extended dark growth **(B)**, respectively. *Orange hue* indicates upregulated proteases, and *blue hue* indicates downregulated proteases. LFQ, label free quantitation.

Proteases were more abundant in LL or DD growth than in the L/D diurnal growth (Figure 4, A, B). This suggests that proteases could play a role in fine-tuning the *Crocosphaera* 51142 proteome in extended periods of light or dark. Among all the proteases identified in LL or DD, an uncharacterized protease (cce_0593) was the most abundant protease followed by the ATP-dependent Clp protease regulatory subunit (ClpC1) (Figure 4B, Table 1). More proteases were upregulated in LL compared to DD. In addition to the ATP dependent Clp proteases (ClpP, ClpP1, ClpP4, ClpX) and carboxy-terminal protease (cce_1372), the zinc metalloproteases (cce_4119, cce_3432, FtsH, FtsH2), protease IV (SppA1 and SppA2), and uncharacterized proteases (cce_4255, cce_4291, cce_2982) were all more abundant in LL compared to DD, whereas processing protease (YmxG) and uncharacterized protease (cce_0593, cce_3195) were more abundant in DD than in LL (Figure 4B, Table 1).

Proteases regulate proteome via protein turnover, and the removal of damaged or inactivated polypeptides [38]. Their functions also extend to recycling of amino acids and turnover of specific, short-lived regulatory proteins that control various metabolic and developmental processes [39]. The ATP-dependent Clp proteases in cyanobacteria, as in many other organisms, form a multigene family, resulting in the production of different isozymes [40]. An 8-fold increase of ClpP1 in *Synechococus* sp. PCC 7942 cells grown at 50 mmol photons m^-2^S^-1^ after supplementing with 0.5 W m^-2^ UV-B for 8 h, suggests that ClpA1 is essential for acclimation to UV-B [41]. A significant increase in the abundances of multiple Clp proteases in *Crocosphaera* 51142 during continuous light exposure in the current investigation indicates that these proteases play an important role in the cellular response of *Crocosphaera* 51142 to adapt to light exposure.

Cyanobacteria express two SppA homologues, SppA1 and SppA2. Although only SppA1 was found to be statistically significant, both SppA1 and SppA2 were more abundant in LL than in DD (Figure 4B, Table 1). The SppA homologues are bacterial integral membrane peptidase [42, 43] and were identified as thylakoid membrane associated proteins in Arabidopsis. This suggested a potential role in processing of the photosynthetic membrane complexes, in addition, their expression and activity are responsive to changes in light conditions [44]. In eukaryotes, the primary ATP-dependent proteolytic system involves the ubiquitin conjugation to target proteins and their subsequent degradation via the multi-subunit proteasome [45]. However, in unicellular cyanobacteria, the exact mechanism by which Clp and other proteases participate in the degradation and recycling of the damaged and inactivated proteins is poorly understood. Our results provide some initial insights into how proteases respond to different light exposure. While the exact mechanisms are currently elusive, these results serve as a foundation for future studies in protein degradation and recycling of damaged or inactivated proteins in unicellular cyanobacteria.

### Ribosomal proteins responsive to different light-dark periods

Low growth rates are often linked to low levels of ribosomal proteins [46]. An interesting observation was that ribosomal proteins were more tightly regulated under L/D than under LL/DD growth (Figure 5A). This may indicate a cell’s ability to maintain metabolic homeostasis under normal light-dark diurnal growth conditions. In contrast, LL, or DD conditions, characterized by continuous light or darkness, respectively, may induce stress on cells due to the absence of a natural light-dark cycle. This stress could lead to dysregulation of translational machinery, resulting in less tightly controlled expression of ribosomal proteins. Calls may struggle to adapt to the constant conditions, leading to a less coordinated response in ribosomal protein regulation. Under L/D, most of the 30S ribosomal proteins showed upregulation during the light period whereas the response of 50S ribosomal proteins to light was mixed (Supplementary Figure 2C). The ribosome recycling factor (Frr, cce_3592) and ribosome hibernation promoting factor (lrtA, cce_0155), which play a role in ribosomal activity, were up in the dark under L/D (Table 1). This variability may be attributed to increased oxidative stress due to extended exposure to continuous light or continuous darkness.

**Figure 5.**
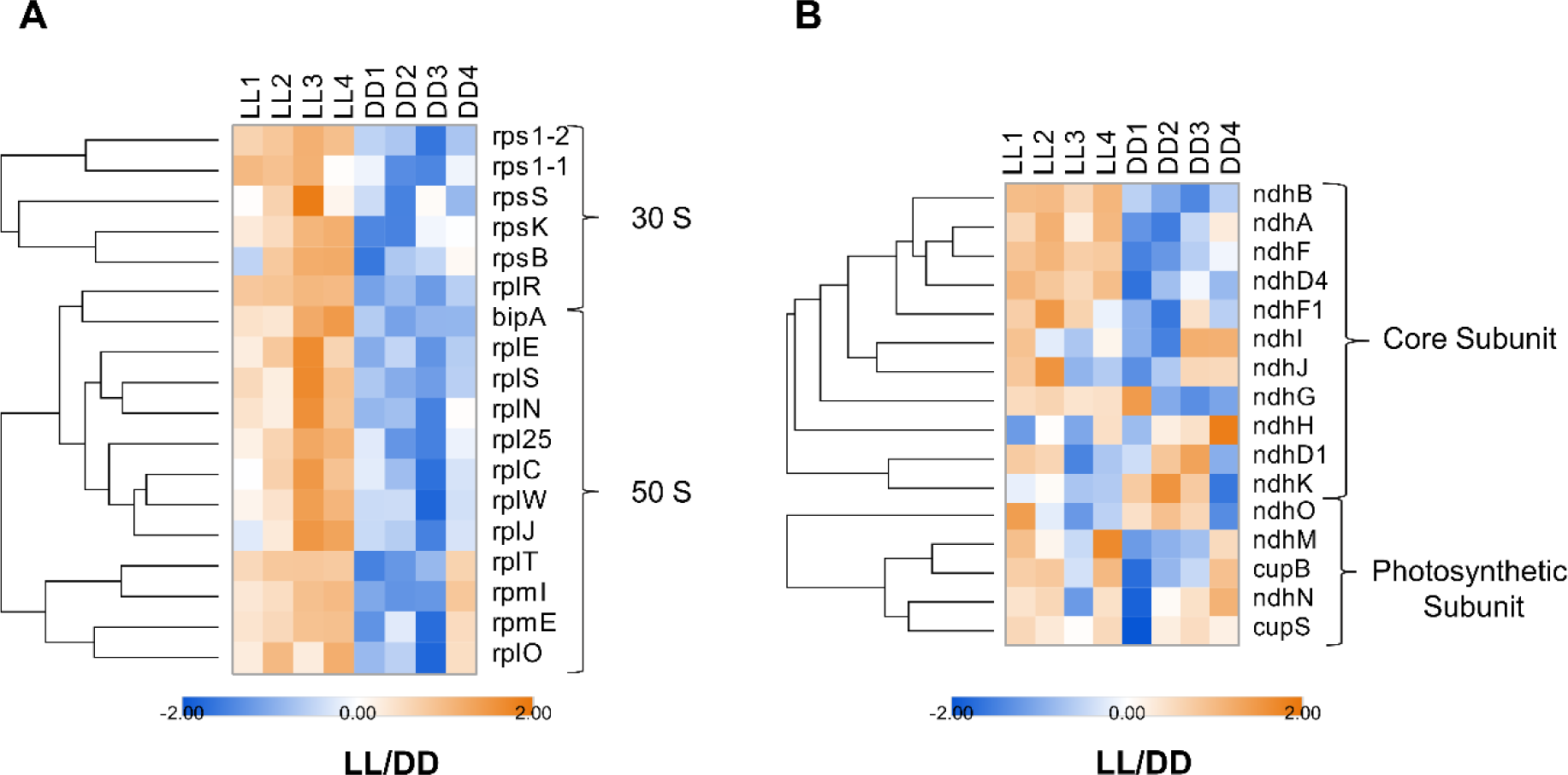
Response of ribosomal and NDH complex subunits. **(A).** Heatmap depicting the z- scored log_2_(LFQ) values of differentially regulated ribosomal protein subunits between extended light and extended dark. Proteins that are a part of 30S and 50S subunits are indicated. **(B).** Heatmap depicting the z-scored log_2_(LFQ) values of differentially regulated NDH complex subunits between extended light and extended dark. Proteins in the homologous core and oxygenic photosynthesis core are indicated. *Orange hue* indicates upregulated proteases, and *blue hue* indicates downregulated proteases. LFQ, label free quantitation.

### NDH complex subunits responsive to light-dark periods

The conversion of light energy into chemical energy during photosynthesis is divided into the “light” and “dark” reactions [47]. The light reaction generates ATP and NADPH which is consumed during the dark reaction to fix CO_2_. The Ndh-1 complex subunits are conserved in cyanobacteria and exhibit functional multiplicity for a variety of functions including respiration, cyclic electron flow, and CO_2_ uptake [48, 49]. They contribute to protecting plants against stress for efficient photosynthesis [50]. We identified both homologous core subunits (Ndh-A, B, D1, D4, F, F1, F4, G, H, I, J, K) as well as oxygenic photosynthetic specific (OPS) subunits (Ndh-L, M, N, O, CupB, and CupS) (Figure 5B, Table 1). However, not all subunits were differentially regulated. Under L/D growth, only two subunits, NdhI and NdhN were upregulated in the dark (Supplementary Figure 2D). The NdhI is associated with Fd-binding and maintenance of FeS cluster in cyanobacteria [47]. Under LL and DD, NdhA, NdhB, NdhD4, NdhF, and NdhF1 were all upregulated under LL (Figure 5B). These core complex subunits are all involved in proton pumping, PQ- and Fd-binding. Cyanobacterial Ndh-1 complexes also participate in active CO_2_ concentration [51]. Upregulation of Ndh-1 complex subunits in the LL suggests their role in protecting cells from oxidative damage and for the maintenance of electron flow and photosynthesis efficiency. Previously, we have shown association between the Cup complexes (NdhN/D3/F3/CupA/CupS complex) to the Ndh-1 core complexes in a the high molecular weight (669 kDa) size exclusion chromatography (SEC) and mass spectrometry analysis and demonstrated the possibility of a functionally diverse Ndh-1 complex including respiration, cyclic electron flow, and CO_2_ uptake form within the same cell at the same time [48].

## DISCUSSION

Changes in protein abundances reflect the changes in cellular strategy to adapt and adjust to changing environmental cues such as light. We identified 603 proteins significantly different between LL/DD growth. The main metabolic enzymes involved in photosynthesis, respiration, co- factors recycling, carbohydrate metabolism, protein translation, and protein degradation showed strong response to light. Although transcript levels of many PSI and PSII proteins were maximal during the day [1, 52], their protein levels were generally more abundant in the night [6-8, 13, 30]. Higher abundances of PSI and PSII proteins in the dark also has been reported previously [6, 13]. This finding may suggest that there is a more rapid protein degradation during the day [13, 30] as evidenced from more prevalent proteases in the light than in the dark (Table 1). This may also relate to the delayed peak of PSI and PSII protein abundances compared to their corresponding transcripts. This adaptation likely ensures that there are sufficient PSI and PSII proteins available for photosynthesis even after the period of peak transcription during the day as there is a more rapid degradation of transcripts and proteins during the light period, as evidenced by increased abundances of several proteases (Figure 4A, Tale 1). Interestingly, this pattern was reversed in cells grown under extended light and dark periods, likely indicating other factors that need to be investigated. One straight forward level of regulation in DD is the downregulation of the luminal protein, which should destabilize the O_2_-evolving complex and lead to lower O_2_ evolution. In addition, the soluble plastocyanin, PetE, is strongly downregulated indicating a decreased need for PSI activity.

One of the interesting observations of this study was the response of different classes of proteases to different light conditions (Figure 4, Table 1). Extreme environments such as excessive light exposure can disturb protein homeostasis (proteostasis) leading to protein misfolding, denaturation, and oxidative damage [53]. Cyanobacteria have evolved protein quality control mechanisms to sense and adapt to such extreme environments [54, 55]. Unlike eukaryotes, which utilize ubiquitin-proteasome system and autophagy, cyanobacteria depend on proteases. This system involves diverse proteolytic machines, notably ATP-dependent Clp proteases [53, 56] and metalloproteases [57]. Significant accumulation of proteases including ClpX, ClpP, FtsH2, and SppA1 (Figure 4B) in LL compared to DD demonstrate the important roles these proteases play for maintaining cellular metabolic activities by disposing and degrading damaged or inactivated proteins [58].

The regulation of proteases in response to light and dark periods suggests intriguing dynamics in protein turnover and cellular processes in *Crocosphaera* 51142 and may reflect distinct metabolic states and cellular activities. Increased abundances of specific proteases in the light could be linked to increased protein turnover and degradation to recycle damaged or misfolded proteins generated during photosynthesis. The CLP proteases, which showed higher abundances in the light are known to play a role in the regulation and maintenance of photosynthetic machinery as certain components of the photosynthetic system may need to be selectively degraded and replaced [59]. The ClpXP complex is a representative proteolytic system in cyanobacteria and consists of hexameric ATPase ClpX, which functions by forming a complex with a tetradecameric peptidase ClpP [60, 61]. In *Synechocystis* sp. PCC6803, disruption of *clpX* led to slower growth, decreased high light tolerance, and impaired photosynthetic cyclic electron transfer. Thus, CLP proteases can mediate these processes, ensuring optimal photosynthetic efficiency and adaptation to fluctuating environment. The regulation of different proteases in the dark, on the other hand, may indicate a shift in cellular priorities, and maybe a lower demand for protein degradation and synthesis in the dark compared to the light. This may result in a relatively stable pool of membrane intrinsic proteins, as their turnover rate tends to be lower compared to soluble proteins [62]. Further investigation into the specific proteases involved and their substrate preferences will provide valuable insights into the mechanism governing protein turnover and cellular adaptation to environmental changes.

A better way to enhance photosynthesis is to improve a cell’s ability to quickly repair impaired PSII systems. When repair cannot match damage, there is a net loss of PSII activity. The FtsH family proteases are membrane-integral ATP-dependent metalloproteases that play a key role in PSII biogenesis and quality control by positively regulating high light inducible proteins [63]. In *Synechocystis* sp. PCC6803, the FtsH2/FtsH3 protease heterocomplex physically interacts with Psb29 subunit and facilitates PSII repair [57]. The downregulation of FtsH results in oxidative stress and the production of reactive oxygen species [57, 64]. The accumulation of FtsH family proteins, including FtsH/FtsH2 in the extended light, may suggest their involvement in proteolysis of damaged PSII complex, including the D1 protein, a core subunit of PSII and the maintenance and acclimation of *Crocosphaera* 51142 cells to extended light exposure. The SppA1 and SppA2, are previously reported light-activated at both the transcriptional and translational levels [44]. In *Synechocystis*, knock-out mutants for each of the SppA1 and SppA2 showed different outputs [65]. The *sppA1* mutant grew faster than the wild type under moderate light regime, whereas the *sppA2* mutant grew slower than the wild type and was not light sensitive. A 5-fold increase in the abundance of SppA1 and 1.6-fold increase of SppA2 in LL compared to DD suggest that SppA1 and its regulation are highly sensitive to light and could play a significant role in light-mediated cellular processes, potentially including stress responses, photosynthetic regulation, or repair mechanisms.

## CONCLUSIONS

In summary, our proteomic results highlight the diverse responses of *Crocosphaera* 51142 cells to extended light and dark conditions. Since these conditions are unanticipated, cells appear to experience stress and adjust such growth conditions by regulating different protein groups. For example, the upregulation of proteins related to photosynthesis and translation during L/D suggests an active phase, while the downregulation of cell wall-related proteins may indicate a reduced emphasis on structural processes and growth. Their distinct response patterns between light and dark periods may indicate that many peptidases identified in this study were not due to functional redundancy as there seems to be little overlap in their regulation in the normal L/D and LL/DD conditions. Despite their importance in a photoprotective role, very little is known about the mechanism underlying these processes in cyanobacteria. Therefore, our results have implications for biotechnological application of photosynthetic cyanobacteria. The results indicate cellular strategy these microbes utilize to minimize protein turnover energy costs and improve fitness under various growth conditions. This could be particularly important for photosynthetic cyanobacteria as their energy supply is limited to times of the day. Furthermore, significant regulation of different classes of proteases in this study may also suggest that changes in transcription, translation, and post-translational events are not sufficient for predicting changes in protein abundances, as various other forces are at play to maintain proteostasis. Further investigations aimed at determining proteases-substrates relationships, the contribution of proteases in protein complex assembly and interactions as well as elucidating functions and mechanisms of these peptidases under different environmental conditions are required. This study could serve as a valuable resource for future studies aimed at understanding the mechanisms governing proteostasis in cyanobacteria, shedding light on their adaptation and survival strategies under varying environmental conditions.

## Data Availability

All the raw LC-MS/MS data are deposited in MassIVE data repository (massive.ucsd.edu) with MASSIVE-ID: MSV000094518

The data for the peptides is available in the MS Viewer repository and can be accessed using the following URL: **https://msviewer.ucsf.edu/prospector/cgi-bin/mssearch.cgi?report_title=MS-Viewer&search_key=zjpgh8mm2y&search_name=msviewer**

The search key for the saved data set is zjpgh8mm2y Click on the URL to display the report.

## Acknowledgements

All LC-MS/MS experiments were performed at the Purdue Proteomics Facility in the Bindley Bioscience Center of Purdue University. We thank Rodrigo Mohallem and other members of the Aryal lab for discussion and feedback in data analysis and interpretation.

## Authors contributions

**Punyatoya Panda**: Culture growth, treatments, proteomics sample preparation, LC-MS/MS data acquisition, data analysis, writing original draft, review, and editing. **Swagarika J. Giri**: GO annotation, data analysis, writing original draft, review, and editing. **Louis Sherman**: conceptualization, project supervision, data interpretation, review, and editing. **Daisuke Kihara**: conceptualization, data analysis, interpretation, manuscript editing, fund acquisition. **Uma K. Aryal**: conceptualization, methodology, data collection, data analysis and interpretation, writing original manuscript, review, and editing, fund acquisition, project supervision.

## Notes

The authors declare no competing financial interests.

## Abbreviations

LC-MS/MS: Liquid Chromatography-Tandem Mass Spectrometry
RSLC: Rapid Separation Liquid Chromatography
HEPES-KOH: 4-(2-hydroxyethyl)piperazine-1-ethanesulfonic acid potassium salt
ATCC: American Type Culture Collection
L: Cells harvested four hours into the light cycle
D: Cells harvested four hours into the dark cycle
LL: Cells harvested 48 hours after continuous light growth
DD: Cells harvested 48 hours after continuous dark growth
BNF: Biological Nitrogen Fixation
PSI: and PSII Photosystem I and Photosystem II
GO: Gene Ontology
CO_2_: Carbon dioxide
N_2_: Dinitrogen
Nif: Nitrogenase enzymes
O_2_: Oxygen
PQC: Protein Quality Control
ATP: Adenosine Triphosphate
AAA+: Protease ATP Associated Protease
NADPH: Nicotinamide Adenine Dinucleotide Phosphate

## Notes

The authors declare no competing financial interests.

## Supporting Information

The following supplementary figures and tables can be accessed free of cost at the ACS website:

**Supplementary Figure 1.** Comparison of proteomic changes during diurnal cycles and extended conditions.

**Supplementary Figure 2.** Responses of photosynthesis and ribosomal proteins.

## Supplementary Tables

**Supplementary Table S1.:** All peptides identified in L,D,LL,DD global analysis

**Supplementary Table S2:** All proteins identified in L,D,LL,DD global analysis

**Supplementary Table S3:** Proteins identified across L,D,LL,DD with at least 70% replicates in at least one group

**Supplementary Table S4:** Two-tailed T-test of L vs D

**Supplementary Table S5:** Two-tailed T-test of LL vs DD

## Index

**Light (L)** Extracted 4 hours into the light cycle

**Dark (D)** Extracted 4 hours into the dark cycle

**Extended Light (LL)** Extracted 48 hours into the extended light cycle

**Extended Dark (DD)** Extracted 48 hours into the extended dark cycle

## References

1. Stockel J, Welsh EA, Liberton M, Kunnvakkam R, Aurora R, Pakrasi HB: Global transcriptomic analysis of Cyanothece 51142 reveals robust diurnal oscillation of central metabolic processes. Proc Natl Acad Sci U S A 2008, 105(16):6156–6161.

2. Welsh EA, Liberton M, Stockel J, Loh T, Elvitigala T, Wang C, Wollam A, Fulton RS, Clifton SW, Jacobs JM et al: The genome of Cyanothece 51142, a unicellular diazotrophic cyanobacterium important in the marine nitrogen cycle. Proc Natl Acad Sci U S A 2008, 105(39):15094–15099.

3. Sherman LA, Meunier P, Colon-Lopez MS: Diurnal rhythms in metabolism: A day in the life of a unicellular, diazotrophic cyanobacterium. Photosynth Res 1998, 58(1):25–42.

4. Schneegurt MA, Sherman DM, Sherman LA: Composition of the carbohydrate granules of the cyanobacterium, Cyanothece sp. strain ATCC 51142. Arch Microbiol 1997, 167(2-3):89-98.

5. Reddy KJ, Haskell JB, Sherman DM, Sherman LA: Unicellular, aerobic nitrogen-fixing cyanobacteria of the genus Cyanothece. J Bacteriol 1993, 175(5):1284–1292.

6. Aryal UK, Callister SJ, Mishra S, Zhang X, Shutthanandan JI, Angel TE, Shukla AK, Monroe ME, Moore RJ, Koppenaal DW et al: Proteome analyses of strains ATCC 51142 and PCC 7822 of the diazotrophic cyanobacterium Cyanothece sp. under culture conditions resulting in enhanced H(2) production. Appl Environ Microbiol 2013, 79(4):1070–1077.

7. Aryal UK, Stockel J, Krovvidi RK, Gritsenko MA, Monroe ME, Moore RJ, Koppenaal DW, Smith RD, Pakrasi HB, Jacobs JM: Dynamic proteomic profiling of a unicellular cyanobacterium Cyanothece ATCC51142 across light-dark diurnal cycles. BMC Syst Biol 2011, 5:194.

8. Aryal UK, Stockel J, Welsh EA, Gritsenko MA, Nicora CD, Koppenaal DW, Smith RD, Pakrasi HB, Jacobs JM: Dynamic proteome analysis of Cyanothece sp. ATCC 51142 under constant light. J Proteome Res 2012, 11(2):609-619.

9. Toepel J, Welsh E, Summerfield TC, Pakrasi HB, Sherman LA: Differential transcriptional analysis of the cyanobacterium Cyanothece sp. strain ATCC 51142 during light-dark and continuous-light growth. J Bacteriol 2008, 190(11):3904–3913.

10. Liberton M, Austin JR, 2nd, Berg RH, Pakrasi HB: Unique thylakoid membrane architecture of a unicellular N2-fixing cyanobacterium revealed by electron tomography. Plant Physiol 2011, 155(4):1656-1666.

11. Elvitigala T, Stockel J, Ghosh BK, Pakrasi HB: Effect of continuous light on diurnal rhythms in Cyanothece sp. ATCC 51142. BMC Genomics 2009, 10:226.

12. Garcia-Dominguez M, Muro-Pastor MI, Reyes JC, Florencio FJ: Light-dependent regulation of cyanobacterial phytochrome expression. J Bacteriol 2000, 182(1):38–44.

13. Stockel J, Jacobs JM, Elvitigala TR, Liberton M, Welsh EA, Polpitiya AD, Gritsenko MA, Nicora CD, Koppenaal DW, Smith RD et al: Diurnal rhythms result in significant changes in the cellular protein complement in the cyanobacterium Cyanothece 51142. PLoS One 2011, 6(2):e16680.

14. Schneegurt MA, Sherman DM, Nayar S, Sherman LA: Oscillating behavior of carbohydrate granule formation and dinitrogen fixation in the cyanobacterium Cyanothece sp. strain ATCC 51142. J Bacteriol 1994, 176(6):1586–1597.

15. Bandyopadhyay A, Elvitigala T, Welsh E, Stockel J, Liberton M, Min H, Sherman LA, Pakrasi HB: Novel metabolic attributes of the genus cyanothece, comprising a group of unicellular nitrogen-fixing Cyanothece. mBio 2011, 2(5).

16. Krishnakumar S, Gaudana SB, Viswanathan GA, Pakrasi HB, Wangikar PP: Rhythm of carbon and nitrogen fixation in unicellular cyanobacteria under turbulent and highly aerobic conditions. Biotechnol Bioeng 2013, 110(9):2371–2379.

17. Bandyopadhyay A, Elvitigala T, Liberton M, Pakrasi HB: Variations in the rhythms of respiration and nitrogen fixation in members of the unicellular diazotrophic cyanobacterial genus Cyanothece. Plant Physiol 2013, 161(3):1334–1346.

18. Kim SQ, Mohallem R, Franco J, Buhman KK, Kim KH, Aryal UK: Multi-Omics Approach Reveals Dysregulation of Protein Phosphorylation Correlated with Lipid Metabolism in Mouse Non-Alcoholic Fatty Liver. Cells 2022, 11(7).

19. Mohallem R, Aryal UK: Regulators of TNFalpha mediated insulin resistance elucidated by quantitative proteomics. Sci Rep 2020, 10(1):20878.

20. Cox J, Mann M: MaxQuant enables high peptide identification rates, individualized p.p.b.-range mass accuracies and proteome-wide protein quantification. Nat Biotechnol 2008, 26(12):1367–1372.

21. Tyanova S, Cox J: Perseus: A Bioinformatics Platform for Integrative Analysis of Proteomics Data in Cancer Research. Methods Mol Biol 2018, 1711:133–148.

22. Tyanova S, Temu T, Sinitcyn P, Carlson A, Hein MY, Geiger T, Mann M, Cox J: The Perseus computational platform for comprehensive analysis of (prote)omics data. Nat Methods 2016, 13(9):731–740.

23. Hawkins T, Luban S, Kihara D: Enhanced automated function prediction using distantly related sequences and contextual association by PFP. Protein Sci 2006, 15(6):1550–1556.

24. Hawkins T, Chitale M, Luban S, Kihara D: PFP: Automated prediction of gene ontology functional annotations with confidence scores using protein sequence data. Proteins 2009, 74(3):566–582.

25. Jain A, Kihara D: Phylo-PFP: improved automated protein function prediction using phylogenetic distance of distantly related sequences. Bioinformatics 2019, 35(5):753–759.

26. Chitale M, Hawkins T, Park C, Kihara D: ESG: extended similarity group method for automated protein function prediction. Bioinformatics 2009, 25(14):1739–1745.

27. Mohallem R, Aryal UK: Quantitative Proteomics and Phosphoproteomics Reveal TNF-alpha-Mediated Protein Functions in Hepatocytes. Molecules 2021, 26(18).

28. Zembroski AS, Buhman KK, Aryal UK: Proteome and phosphoproteome characterization of liver in the postprandial state from diet-induced obese and lean mice. J Proteomics 2021, 232:104072.

29. Shen JR, Qian M, Inoue Y, Burnap RL: Functional characterization of Synechocystis sp. PCC 6803 delta psbU and delta psbV mutants reveals important roles of cytochrome c-550 in cyanobacterial oxygen evolution. Biochemistry 1998, 37(6):1551-1558.

30. Sherman LA, Meunier P, Colón-López MS: Diurnal rhythms in metabolism:: A day in the life of a unicellular, diazotrophic cyanobacterium. Photosynth Res 1998, 58(1):25–42.

31. Gaudana SB, Alagesan S, Chetty M, Wangikar PP: Diurnal rhythm of a unicellular diazotrophic cyanobacterium under mixotrophic conditions and elevated carbon dioxide. Photosynth Res 2013, 118(1-2):51–57.

32. Kohler U, Cerff R, Brinkmann H: Transaldolase genes from the cyanobacteria Anabaena variabilis and Synechocystis sp. PCC 6803: comparison with other eubacterial and eukaryotic homologues. *Plant Mol Biol* 1996, 30(1):213-218.

33. Lee JZ, Burow LC, Woebken D, Everroad RC, Kubo MD, Spormann AM, Weber PK, Pett-Ridge J, Bebout BM, Hoehler TM: Fermentation couples Chloroflexi and sulfate-reducing bacteria to Cyanobacteria in hypersaline microbial mats. Front Microbiol 2014, 5:61.

34. Sherman LA, Min H, Toepel J, Pakrasi HB: Better living through cyanothece - unicellular diazotrophic cyanobacteria with highly versatile metabolic systems. Adv Exp Med Biol 2010, 675:275–290.

35. Liberton M, Bandyopadhyay A, Pakrasi HB: Enhanced Nitrogen Fixation in a glgX- Deficient Strain of Cyanothece sp. Strain ATCC 51142, a Unicellular Nitrogen-Fixing Cyanobacterium. Appl Environ Microbiol 2019, 85(7).

36. Suzuki E, Umeda K, Nihei S, Moriya K, Ohkawa H, Fujiwara S, Tsuzuki M, Nakamura Y: Role of the GlgX protein in glycogen metabolism of the cyanobacterium, Synechococcus elongatus PCC 7942. Biochim Biophys Acta 2007, 1770(5):763-773.

37. Bouchnak I, van Wijk KJ: Structure, function, and substrates of Clp AAA+ protease systems in cyanobacteria, plastids, and apicoplasts: A comparative analysis. J Biol Chem 2021, 296:100338.

38. Burgos R, Weber M, Martinez S, Lluch-Senar M, Serrano L: Protein quality control and regulated proteolysis in the genome-reduced organism Mycoplasma pneumoniae. Mol Syst Biol 2020, 16(12):e9530.

39. Parsell DA, Lindquist S: The function of heat-shock proteins in stress tolerance: degradation and reactivation of damaged proteins. Annu Rev Genet 1993, 27:437–496.

40. Schelin J, Lindmark F, Clarke AK: The clpP multigene family for the ATP-dependent Clp protease in the cyanobacterium Synechococcus. Microbiology (Reading) 2002, 148(Pt 7):2255–2265.

41. Porankiewicz J, Schelin J, Clarke AK: The ATP-dependent Clp protease is essential for acclimation to UV-B and low temperature in the cyanobacterium Synechococcus. Mol Microbiol 1998, 29(1):275–283.

42. Bolhuis A, Matzen A, Hyyrylainen HL, Kontinen VP, Meima R, Chapuis J, Venema G, Bron S, Freudl R, van Dijl JM: Signal peptide peptidase- and ClpP-like proteins of Bacillus subtilis required for efficient translocation and processing of secretory proteins. J Biol Chem 1999, 274(35):24585–24592.

43. Novak P, Dev IK: Degradation of a signal peptide by protease IV and oligopeptidase A. J Bacteriol 1988, 170(11):5067–5075.

44. Lensch M, Herrmann RG, Sokolenko A: Identification and characterization of SppA, a novel light-inducible chloroplast protease complex associated with thylakoid membranes. J Biol Chem 2001, 276(36):33645–33651.

45. Rechsteiner M: Ubiquitin-mediated pathways for intracellular proteolysis. Annu Rev Cell Biol 1987, 3:1-30.

46. Luimstra VM, Schuurmans JM, Hellingwerf KJ, Matthijs HCP, Huisman J: Blue light induces major changes in the gene expression profile of the cyanobacterium Synechocystis sp. PCC 6803. *Physiol Plant* 2020, 170(1):10-26.

47. Laughlin TG, Savage DF, Davies KM: Recent advances on the structure and function of NDH-1: The complex I of oxygenic photosynthesis. Biochim Biophys Acta Bioenerg 2020, 1861(11):148254.

48. Aryal UK, Ding Z, Hedrick V, Sobreira TJP, Kihara D, Sherman LA: Analysis of Protein Complexes in the Unicellular Cyanobacterium Cyanothece ATCC 51142. J Proteome Res 2018, 17(11):3628–3643.

49. Miller NT, Vaughn MD, Burnap RL: Electron flow through NDH-1 complexes is the major driver of cyclic electron flow-dependent proton pumping in cyanobacteria. Biochim Biophys Acta Bioenerg 2021, 1862(3):148354.

50. Hualing M: Cyanobacterial NDH-1 Complexes. Front Microbiol 2022, 13:933160.

51. Klughammer B, Sultemeyer D, Badger MR, Price GD: The involvement of NAD(P)H dehydrogenase subunits, NdhD3 and NdhF3, in high-affinity CO2 uptake in Synechococcus sp. PCC7002 gives evidence for multiple NDH-1 complexes with specific roles in cyanobacteria. Mol Microbiol 1999, 32(6):1305-1315.

52. Colon-Lopez MS, Sherman LA: Transcriptional and translational regulation of photosystem I and II genes in light-dark- and continuous-light-grown cultures of the unicellular cyanobacterium Cyanothece sp. strain ATCC 51142. J Bacteriol 1998, 180(3):519–526.

53. Zhang Y, Wang Y, Wei W, Wang M, Jia S, Yang M, Ge F: Proteomic analysis of the regulatory networks of ClpX in a model cyanobacterium Synechocystis sp. PCC 6803. Front Plant Sci 2022, 13:994056.

54. Franklin DJ: Examining the Evidence for Regulated and Programmed Cell Death in Cyanobacteria. How Significant Are Different Forms of Cell Death in Cyanobacteria Population Dynamics? Front Microbiol 2021, 12:633954.

55. Cui J, Xie Y, Sun T, Chen L, Zhang W: Deciphering and engineering photosynthetic cyanobacteria for heavy metal bioremediation. Sci Total Environ 2021, 761:144111.

56. He Z, Mi H: Functional Characterization of the Subunits N, H, J, and O of the NAD(P)H Dehydrogenase Complexes in Synechocystis sp. Strain PCC 6803. Plant Physiol 2016, 171(2):1320-1332.

57. Bec Kova M, Yu J, Krynicka V, Kozlo A, Shao S, Konik P, Komenda J, Murray JW, Nixon PJ: Structure of Psb29/Thf1 and its association with the FtsH protease complex involved in photosystem II repair in cyanobacteria. Philos Trans R Soc Lond B Biol Sci 2017, 372(1730).

58. Sauer RT, Bolon DN, Burton BM, Burton RE, Flynn JM, Grant RA, Hersch GL, Joshi SA, Kenniston JA, Levchenko I et al: Sculpting the proteome with AAA(+) proteases and disassembly machines. Cell 2004, 119(1):9–18.

59. Nishimura K, Kato Y, Sakamoto W: Chloroplast Proteases: Updates on Proteolysis within and across Suborganellar Compartments. Plant Physiol 2016, 171(4):2280–2293.

60. Glynn SE, Martin A, Nager AR, Baker TA, Sauer RT: Structures of asymmetric ClpX hexamers reveal nucleotide-dependent motions in a AAA+ protein-unfolding machine. Cell 2009, 139(4):744–756.

61. Alexopoulos JA, Guarne A, Ortega J: ClpP: a structurally dynamic protease regulated by AAA+ proteins. J Struct Biol 2012, 179(2):202–210.

62. Karlsen J, Asplund-Samuelsson J, Jahn M, Vitay D, Hudson EP: Slow Protein Turnover Explains Limited Protein-Level Response to Diurnal Transcriptional Oscillations in Cyanobacteria. Front Microbiol 2021, 12:657379.

63. Krynicka V, Skotnicova P, Jackson PJ, Barnett S, Yu J, Wysocka A, Kana R, Dickman MJ, Nixon PJ, Hunter CN et al: FtsH4 protease controls biogenesis of the PSII complex by dual regulation of high light-inducible proteins. Plant Commun 2023, 4(1):100502.

64. Krynicka V, Tichy M, Krafl J, Yu J, Kana R, Boehm M, Nixon PJ, Komenda J: Two essential FtsH proteases control the level of the Fur repressor during iron deficiency in the cyanobacterium Synechocystis sp. PCC 6803. Mol Microbiol 2014, 94(3):609-624.

65. Sokolenko A, Pojidaeva E, Zinchenko V, Panichkin V, Glaser VM, Herrmann RG, Shestakov SV: The gene complement for proteolysis in the cyanobacterium Synechocystis sp. PCC 6803 and Arabidopsis thaliana chloroplasts. Curr Genet 2002, 41(5):291-310.

