## Supporting information for "Proteomic analysis of unicellular cyanobacterium *Crocosphaera subtropica* ATCC 51142 under extended light or dark growth"

**Table of Contents**

**Supplementary Figures**

**Supplementary Figure 1.** Comparison of proteomic changes during diurnal cycles and extended conditions.
**Supplementary Figure 2.** Responses of photosynthesis and ribosomal proteins.
**Supplementary Tables:**
**Supplementary Table S1 :** All peptides identified in L,D,LL,DD global analysis
**Supplementary Table S2:** All proteins identified in L,D,LL,DD global analysis
**Supplementary Table S3:** Proteins identified across L,D,LL,DD with at least 70% replicates in at least one group
**Supplementary Table S4:** Two-tailed T-test of L vs D
**Supplementary Table S5:** Two-tailed T-test of LL vs DD

**Index**

| | |
|---|---|
| **Light (L)** | Extracted 4 hours into the light cycle |
| **Dark (D)** | Extracted 4 hours into the dark cycle |
| **Extended Light (LL)** | Extracted 48 hours into the extended light cycle |
| **Extended Dark (DD)** | Extracted 48 hours into the extended dark cycle |

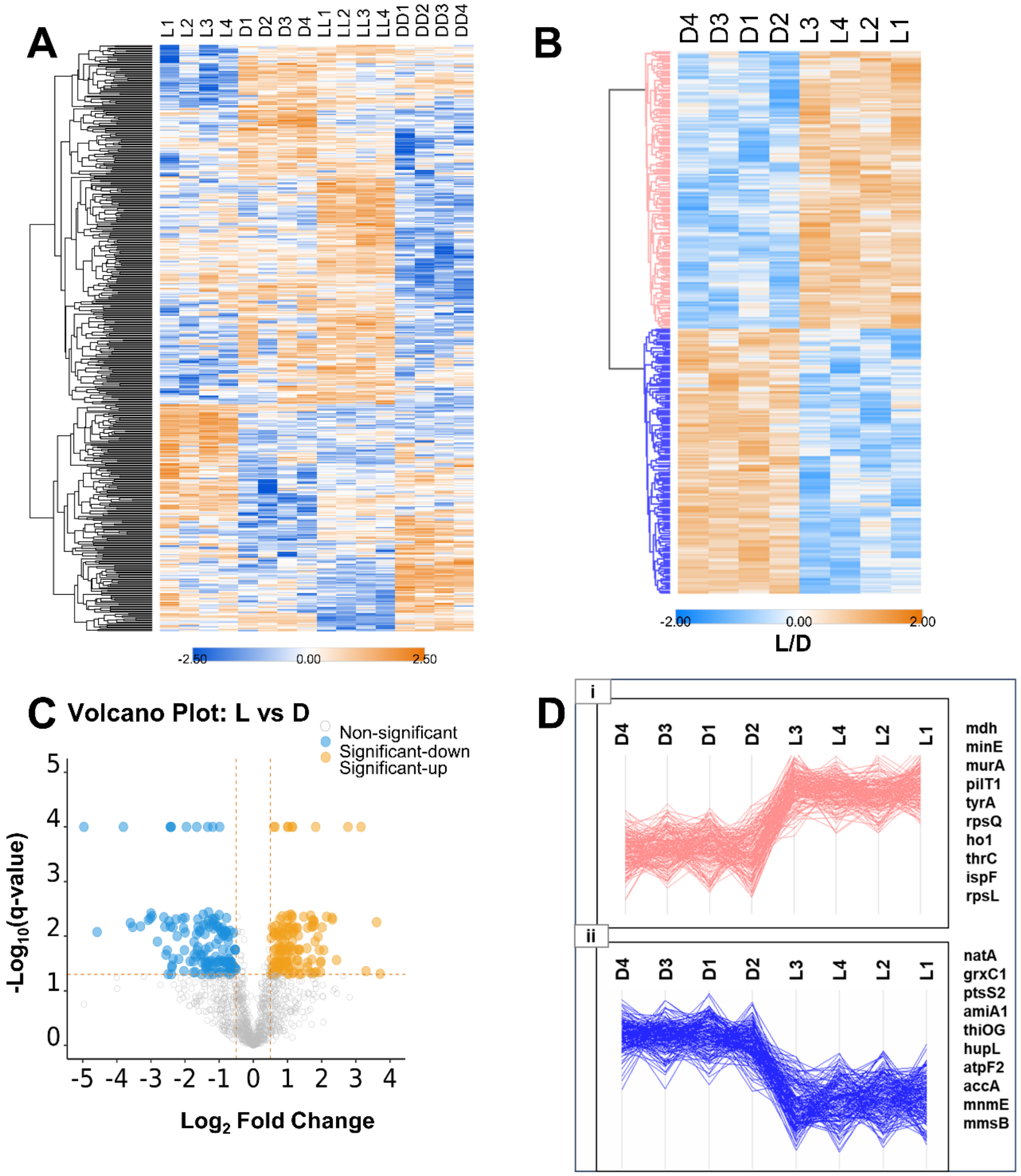


**Supplementary Figure 1. Comparison of proteomic changes during diurnal cycles and extended conditions. (A).** The heatmap of all 1640 proteins identified across light (L), dark (D), extended light (LL) and extended dark (DD) samples. **(B).** Heatmap of 315 significantly changing proteins between light (L) and dark (D). Data represents the z-scored $\log_2$(LFQ) values of all significantly different proteins. LFQ, label free quantitation. **(C).** Volcano plot of all quantified proteins under light and dark conditions. The horizontal line represents the *$\log_2$ (fold change)* cutoff*, and the* vertical line represents the *-$\log_{10}$ (q-value)* cutoff. **(D).** Cluster profiles of significant proteins. The top 10 proteins with the highest and lowest fold change in L and D conditions are indicated on the right side of Figure D.

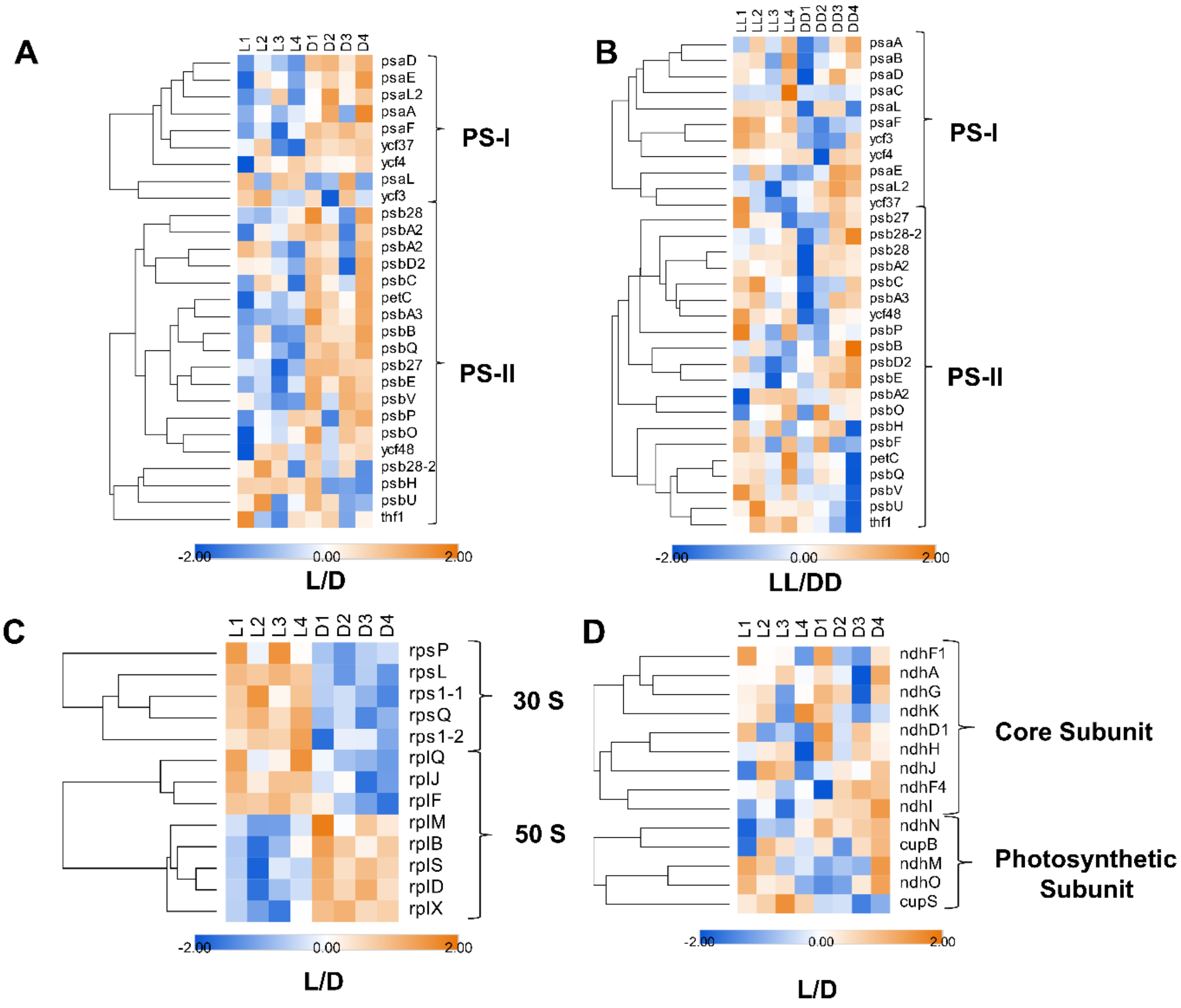


**Supplementary Figure 2. Responses of photosynthesis and ribosomal proteins. (A, B).** Heatmap depicting the z-scored $\log_2$(LFQ) values of photosystem I and II proteins exhibiting differences in abundances between L/D and LL/DD, respectively. **(C, D).** Heatmap depicting the z-scored $\log_2$(LFQ) values of ribosomal protein subunits and NDH-1 complex subunits between light and dark growth, respectively. LFQ, label free quantitation.
